# The Role of Interspecies recombinations in the evolution of antibiotic-resistant pneumococci

**DOI:** 10.1101/2021.02.22.432219

**Authors:** Joshua C. D’Aeth, Mark P.G. van der Linden, Lesley McGee, Herminia De Lencastre, Paul Turner, Jae-Hoon Song, Stephanie W. Lo, Rebecca A. Gladstone, Raquel Sá-Leão, Kwan Soo Ko, William P. Hanage, Bernard Beall, Stephen D. Bentley, Nicholas J. Croucher, The GPS Consortium

## Abstract

The evolutionary histories of the antibiotic-resistant *Streptococcus pneumoniae* lineages PMEN3 and PMEN9 were reconstructed using global collections of genomes. In PMEN3, one resistant clade spread worldwide, and underwent 25 serotype switches, enabling evasion of vaccine-induced immunity. In PMEN9, only 9 switches were detected, and multiple resistant lineages emerged independently and circulated locally. In Germany, PMEN9’s expansion correlated significantly with the macrolide:penicillin consumption ratio. These isolates were penicillin sensitive but macrolide resistant, through a homologous recombination that integrated Tn*1207.1* into a competence gene, preventing further diversification via transformation. Analysis of a species-wide dataset found 183 acquisitions of macrolide resistance, and multiple gains of the tetracycline-resistant transposon Tn*916*, through homologous recombination, often originating in other streptococcal species. Consequently, antibiotic selection preserves atypical recombination events that cause sequence divergence and structural variation throughout the *S. pneumoniae* chromosome. These events reveal the genetic exchanges between species normally counter-selected until perturbed by clinical interventions.

## Introduction

Infections caused by *Streptococcus pneumoniae* (the pneumococcus) remain a leading cause of death worldwide in children under the age of 5 [1,2]. This nasopharyngeal commensal and respiratory pathogen causes a range of severe infections in both infants and adults, including pneumonia, sepsis and meningitis. These have a high mortality rate, which is further increased when the causative pneumococcus is resistant to antibiotics [3, 4]. This presents a worrying challenge to clinicians, with treatment options decreasing for resistant infections [5]. As the pneumococcus is endemic worldwide, its ability to develop antibiotic resistance is a global challenge [6]. Furthermore, this resistance is under selection by community antibiotic consumption, which is driven by common non-invasive pneumococcal diseases, such as otitis media [7, 8]. High levels of resistance have been observed in Africa [9], Asia and the Americas. Even in Europe, where resistance is less common, deaths attributable to penicillin resistant pneumococci have been rising over the past 15 years [3].

There are two main mechanisms by which pneumococci evolve antibiotic resistance: the modification of core genes encoding antibiotic targets through homologous recombination, and the acquisition of specialized resistance genes on mobile genetic elements (MGEs) [7, 10]. As the plasmid repertoire of *S. pneumoniae* is limited to two types of cryptic elements [11–13], the MGEs that contribute most to the spread of antibiotic resistance are integrative and conjugative elements (ICEs) [14, 15]. These MGEs insert within the host genome via encoded integrase genes [16], and have been referred to as king makers of bacterial lineages [17]. Conjugation is a highly efficient method of DNA transfer, with the pilus formed between donor and recipient cells protecting transferred DNA from the external environment [18]. This has enabled conjugation to transfer elements across a broad range of bacterial taxa [19–21], which seems the most likely explanation as to how antibiotic resistance genes originally entered the *S. pneumoniae* population.

The most important ICE driving the spread of antibiotic resistance in pneumococci is Tn*916*, which was the first ICE to be discovered [22, 23]. This element confers tetracycline resistance via the *tetM* gene, and also forms composite elements that can confer resistance to macrolides, amingoglycosides, streptogramins and lincosmides through the integration of sequences such as the Mega cassette, Omega cassette and Tn*917* elements [24]. Tn*916*-type elements are present in antibiotic-resistant pneumococci, and also appear in the majority of antibiotic-resistant bacterial pathogens considered a priority by the WHO [20, 23, 25]. However, the distribution of Tn*916* in *S. pneumoniae* is a particular puzzle, as *in vitro* studies have shown it appears unable to conjugate between pneumococci, although pneumococci themselves can be donors to other streptococci [26,27]. The short cassettes that integrate into Tn*916* also lack their own self-mobilisation machinery, which is even true for the longer form of the Mega cassette, Tn*1207.1*. Hence the contribution of conjugation to the spread of these ICE is not clear.

An alternative mechanism by which Tn*916* might spread is transformation, which facilitates the uptake of extra-cellular DNA into cells which have reached a competent state [28]. Originally discovered in the pneumococcus, transformation is tightly controlled by the host cell, which encodes all the required machinery [28]. Imported DNA is then integrated into the chromosome via homologous recombination. However, there are two important limitations on the dissemination of MGEs between species through transformation.

The first is the inhibition of homologous recombination by sequence divergence, which limits the integration of sequence from other species into host chromosomal DNA [29–32]. The decline is exponential as sequence divergence between the donor and recipient increases [33]. This is thought to result from the minimum effective processing segment (MEPS), the shortest length of continuous sequence identity required for efficient recombination, estimated to be 27 bp for pneumococci [32]. This barrier though, was not sufficient to prevent interspecies recombinations facilitating the emergence of beta lactam resistant pneumococci. This involved the formation of ‘mosaic’ versions of multiple genes encoding targets of beta lactam antibiotics (most commonly, *pbp1a*, *pbp2b* and *pbp2x*) that were a mixture of sequence from *S. pneumoniae* and the related oronasopharyngeal commensal streptococcal species, *Streptococcus mitis* and *Streptococcus oralis* [19, 34, 35]. The mosaicism reflected the imported fragments being much smaller than a typical gene, although there was also evidence of these recombinations causing diversification in the flanking regions of the chromosome [36].

The second limitation to the acquisition of an MGE through transformation, is that transformation requires the importation of both the intact locus and two flanking “homologous arms”, in which recombination crossovers can occur. Both arms must match the host chromosome, and no cleavage of the imported DNA must occur between them during its uptake into the cell. Hence the efficiency of uptake declines with the length of the inserted locus between the two arms [37]. Therefore, while *in vitro* studies have shown that interspecies transfer of MGEs via transformation is possible [38], these transfers were of a low frequency, and only spread shorter resistance cassettes. As such, transformation is thought to spread MGEs primarily through intraspecies transfer of already successfully inserted elements [39].

Therefore, although there are multiple mechanisms by which the MGEs may be acquired, their relative contributions are not known. This is particularly challenging when antibiotic resistance loci are ubiquitous throughout a strain, as Tn*916* is within many AMR *S. pneumoniae* lineages [10, 24, 40], making the process underlying the MGE’s acquisition difficult to infer. Therefore we investigated two globally distributed pneumococcal lineages in which antibiotic resistance MGEs are common, but not conserved.

The first is the Spain^9*V*^ -3, or PMEN3 lineage, which is within strain GPSC6 (clonal complex 156 by multi-locus sequence typing, MLST) [41]. PMEN3 was first documented in Spain in 1988 with a serotype 9V capsule. It was later detected in France, the USA and South America [42–45]. By 2000, 55% of all penicillin resistant disease isolates in South America were from the PMEN3 lineage [45].

The second is the England^14^-9, or PMEN9 lineage, which is within strain GPSC18 (clonal complex 9 or 15 by MLST). This was first described in the UK in 1996 and it has been isolated across Europe, the Americas and Asia [46–48]. PMEN9 was the most common lineage causing penicillin-resistant invasive pneumococcal disease (IPD) in the USA, and the most common lineage causing macrolide-resistant IPD in Germany, prior to the introduction of infant vaccination [49–51].

Both lineages exhibit variability in their resistance to tetracyclines and macrolides across countries, suggesting frequent acquisition or loss of elements such as Tn*916* and Tn*1207.1* [52–55]. Therefore we analysed the distribution of antibiotic resistance loci within these lineages, and then expanded the study across the species with the wider Global Pneumococcal Sequencing project (GPS) data [41, 56]. Using a mixture of genomic approaches, we assessed the distribution of antibiotic resistance loci, determined the mechanisms by which they were imported into these lineages, and characterized how these genotypes have adapted to local antibiotic use as they have spread globally.

## Methods

### Bacterial isolates and DNA sequencing

For the *S. pneumoniae* PMEN3 and PMEN9 datasets, isolates were collated across Europe (from the Nationales Referenzzentrum für Streptokokken, Germany; and the collections of Prof. de Lencastre), the Americas (from the collections of the CDC through the Global Strain Bank Project) and the Maela refugee camp in Thailand [57] (Table S1). These datasets had multi-locus sequence typing (MLST) data [58]. Therefore isolates of sequence type (ST) 156, and single locus variants thereof, could be selected as representatives of PMEN3 (or Spain^9*V*^ -3); isolates of ST9, and single locus variants thereof, were selected as representatives of PMEN9 (or England^14^-9) [59]. This generated collections of 272 and 325 isolates for PMEN3 and PMEN9, respectively. Isolates that could be cultured were sequenced as paired-end 24-plex libraries on Illumina HiSeq 2000 machines, generating 75 nt reads. After comparing inferred serotypes from seroba v1.0.0 [60] and STs to those determined by the sample providers as described previously [24], and manually checking for signals of contamination, 215 PMEN3, and 263 PMEN9, isolates were used in the described analyses. All data were submitted to the ENA, and accession codes are listed in Table S1.

These datasets were combined with isolates from the Global Pneumococcal Sequencing (GPS) project, which generated a database of 20,015 high-quality pneumococcal draft genome sequences from 33 countries collected between 1991 and 2017 [41]. In total, 49.7% of these isolates were collected from locations before the PCV7 vaccine was introduced. The majority of isolates were sampled from cases of invasive pneumococcal disease (IPD) in children under the age of five. PMEN3 and PMEN9 corresponded to strains GPSC6 (454 isolates) and GPSC18 (312 Isolates) in this collection, respectively [61]. Hence the final dataset sizes were 669 for PMEN3 and 575 for PMEN9. Among these PMEN collections, 64.7% of isolates were collected before the PCV7 vaccine was introduced. WGS data from the 478 isolates not within the GPS collection are publically available in the EMBL Nucleotide Sequence Database (ENA; Project number PRJEB2255).

### Generation of annotation and alignments

De novo assemblies were generated using an automated pipeline for Illumina sequences [62]. Briefly, reads were assembled using Velvet with parameters selected by VelvetOptimiser. These draft assemblies were then improved by using SSPACE and GapFiller to join contigs [63–65]. The final assemblies were annotated using PROKKA [66].

Whole genome alignments were generated for phylogenetic analysis through mapping of short read data against reference sequences. For the PMEN3 and PMEN9 analyses, the reference genomes were *S. pneumoniae* RMV4 *rpsL*\* Δ*tvrR* (accession code: ERS1681526) [67] and INV200 (accession code: FQ312029.1) respectively. Mapping was performed using SMALT v0.64, the GATK indel alignment toolkit and SAMtools as described previously [32]. A more computationally-efficient approach was applied to the GPSCs defined from the GPS project. GPSCs with > 10 isolates present, were then aligned to a reference sequence. The reference was chosen as the isolate with the largest N_50_ value (the length of the contig at the midpoint of the assembly, when contigs are ordered by size). Other isolates were mapped to this reference using SKA [68].

### Antibiotic consumption data

Selection pressures on specific clades was estimated using antibiotic consumption data from Europe. Consumption data for macrolides and penicillins was collected. Two sources were used for macrolides, a study looking at macrolide resistance among pneumococci isolates in Germany by Reinert *et al* 2002 [69], which has data from 1992 to 2000, and the European Centre for Disease prevention and Control (ECDC), which has data from 1997 to the present day. The macrolide usage data were combined using the three years of overlap between the two datasets as a scaling factor. This was the average transformation that mapped the Reinert *et al* 2002 data to the ECDC data. It was applied to convert the data from 1992 to 1996 into the same units as the ECDC data (defined daily doses / 1000 population). The ECDC data for Germany is from the primary care sector for outpatients, with a population coverage of 90%, while the Reinert *et al* paper takes data from both prescriptions in hospitals and from community general practitioners.

For penicillin consumption, data was also taken from the ECDC for 1997 to present day. For penicillin consumption from 1992 to 1997 data was taken from McManus *et al* 1997 [70]. This has data on the hospital and retail sales of oral antibiotics in West Germany for the years 1989 and 1994 in the same DDD units as the ECDC data. A linear trend between 1989, 1994 and 1997, the first year of data from the ECDC, was used to impute the missing values between 1992 and 1997.

### Phylogenetic and phylodynamic analyses

Gubbins v2.3 was used to identify recombinations and generate phylogenies for both the PMEN lineages and the GPSCs [71]. This was run for five iterations, with the first phylogeny constructed with FastTree 2 [72], and subsequent iterations generating phylogenies with RAxML v8.2.8 [73], with a generalized time reversible (GTR) model of nucleotide substitution with a discretised gamma distribution of rates across sites.

Time-calibrated phylogenies were generated from the Gubbins outputs using the BactDating R package v1.0.1 [74]. Isolates without dates of collection were pruned from the phylogeny, and the roottotip distances used to test for a molecular clock signal. Where one was detectable, BactDating was run with a relaxed clock model and a Markov chain Monte Carlo (MCMC) length of between 50 and 100 million iterations. Chain convergence was checked through visual inspection of trace plots.

The Skygrowth R package [75] was then used to formally test the link between antibiotic consumption and population growth rates. The timed phylogeny generated by BactDating and the amalgamated macrolide consumption data were input into Skygrowth, with the MCMC run for 50 million iterations for analysis both including and omitting the macrolide usage data.

### Antibiotic resistance analyses

The minimum inhibitory concentration (MIC) for penicillin had been determined for the majority of isolates in the PMEN3 and PMEN9 collections (65.5% and 80.9% respectively). The MICs of the remaining 341 isolates were predicted using a random forest (RF) protocol developed in Li *et al* 2017 [76]. In this protocol, the Transpeptidase domains (TPD) of three penicillin binding proteins (PBPs; PBP1A, PBP2B & PBP2X) were extracted and each amino acid position used as a predictor to train an RF model on the continuous log_2_ MIC value. The training data came from 4,342 isolates previously characterised by the CDC [77].

This model predicted the MIC for 340 of the 341 isolates with unknown MIC values, with the missing isolate having unidentifiable *pbp* genes. The continuous MIC values predicted by the model were then converted into categories based on the pre-2008 meningitis breakpoints for resistance, with an added intermediate class of isolates for those with 0.06 *µ*g/l < MIC < 0.12 *µ*g/l [78].

This method was then used on the wider 20,015 GPS collection, with 59 isolates (0.3%) having unidentifiable *pbp* genes.

Resistance to sulfamethoxazole was detected using a hidden Markov model (HMM), constructed using HMMer3 [79], trained to extract the region downsteam of S61 in *folP*. If this region contained at least one inserted amino acid, then the isolate was predicted to be resistant. Resistance to trimethoprim used an HMM to identify the amino acid at position 100 in *dhfR* (also known as *dyr* or *folk*). Isolates with an isoleucine at this position were predicted to be sensitive, and isolates with a leucine at this position were assumed to be resistant. If isolates were classified as resistant to both sulfamethoxazole and trimethoprim, they were also classified as resistant to the combination drug cotrimoxazole [80].

The code for both the penicillin prediction and co-trimoxazole prediction methods is available at https://github.com/jdaeth274/pbp_tpd_extraction.

### Ancestral state reconstruction

The penicillin resistance categories inferred from the metadata and the RF model were reconstructed on the timecalibrated phylogeny using the phytools R package v0.7.7 [81]. The make.simmap function was run using an equal rates model and an MCMC chain sampling every 100 iterations. The input was a matrix of character states for the tips as probabilities if the phenotype was inferred from the RF model; otherwise, observed phenotypes were assigned a probability of one. For the single tip with no data, the probabilities of each of the states was estimated from their proportions in the rest of the dataset.

Each node’s state was assigned as that with the highest posterior probability. Starting at the root, the number of lineages of each state at each coalescent event in the time-calibrated tree was recorded. Every time a node was reached an extra lineage was added to the total. If there was no state change between two nodes the count for the state was increased by one; else if there was a state change the count for the new state was increased by two, given the strictly bifurcating nature of the timecalibrated tree. The total number and the proportion of branches in each state were recorded through time.

To assess the number of serotype switching events across the PMEN lineages, The JOINT Maximum Likelihood (ML) model for ancestral reconstruction of PastML v1.9.15 [82], was used.

### MGE identification

Reference MGE sequences were used to search the PMEN3, PMEN9 and GPSCs for intact and partial representatives. For the Tn*916* element, the 18 kb reference given by the transposon registry [83], extracted from *Bacillus subtilis* (accession code: KM516885), was used. For Tn*1207.1*, a 7 kb reference extracted from the *S. pneumoniae* INV200 genome (accession code: FQ312029.1) was used. BLASTN was used to detect Tn*916* and Tn*1207.1* among the assembled genomes in the collections, setting an empirically determined cutoff alignment length of 7 kb and 2 kb respectively for each element. BLASTN results were merged if they represented continuation of an element’s sequence split across multiple contigs, to enable detection of elements in isolates that were fragmented in draft assemblies.

### MGE insertion site identification

A pipeline was developed to categorise the insertion points of the elements and infer the node within a cluster’s phylogeny at which the insertion occurred. Figure (1) outlines the algorithm. The initial step was the creation of a library of unique hits, with BLAST matches against the GPSC’s reference determining the start and end points of an insertion. A hit was defined by three charecteristics: (1) the total length of the delineated insertion; (2) the number of genes within the insertion, and (3) the genes within the flanking regions of a hit. Each observed combination of values was considered a unique hit. For instance, if two hits were of similar length and gene content, but differ in where they insert within the host, they were treated as two unique hits. The unique insertions with the largest flanking matches to the reference, indicating the insert was reconstructed on a large contig, were used as representatives of that insertion within the library.

**Figure 1:**
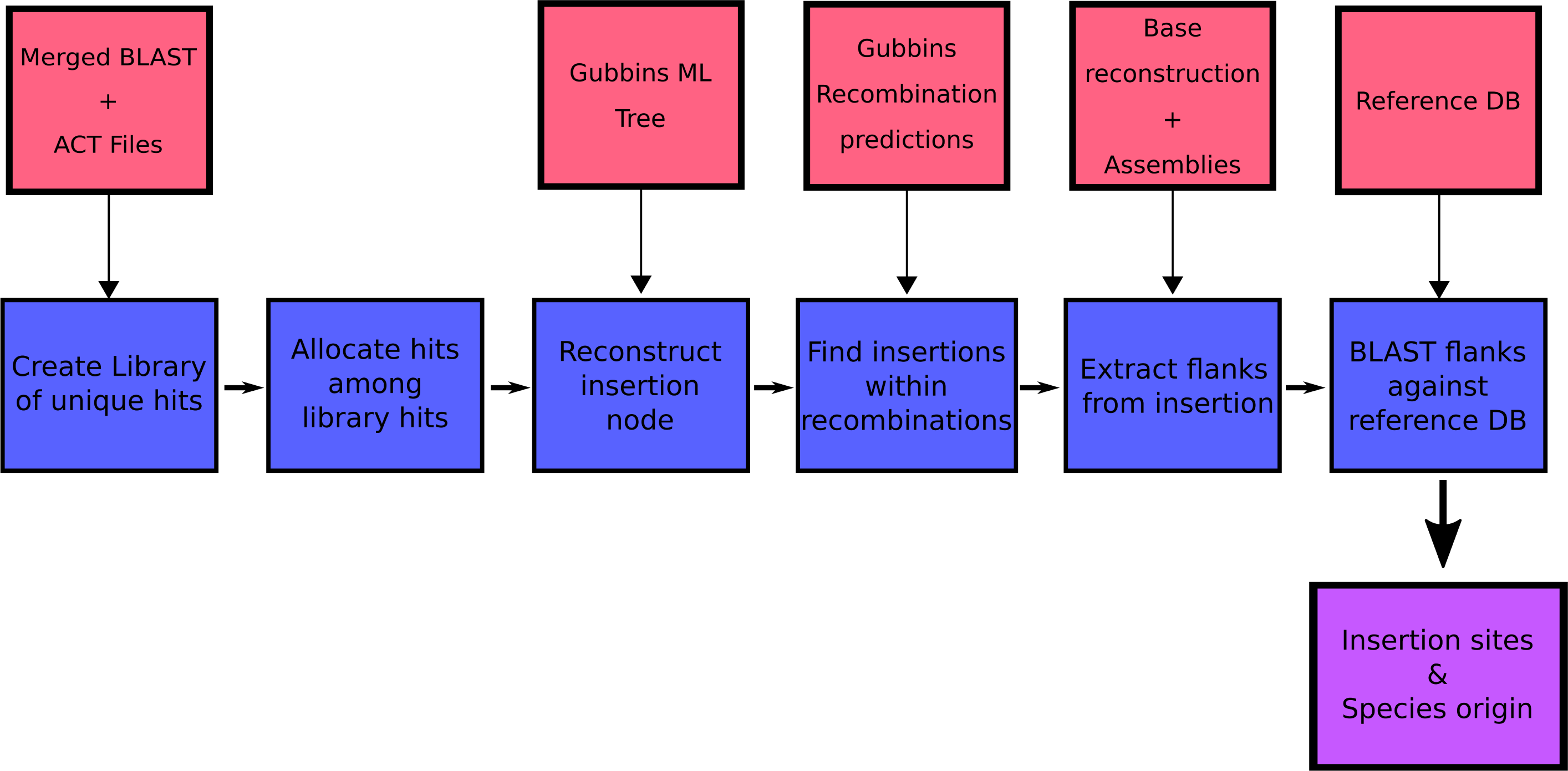
Outline of insertion point pipeline. Red boxes represent data input into the pipeline, blue boxes the individual analysis steps within the pipeline and purple the pipeline’s output.

The next step was to allocate the remaining hits, which were not present in the library, to one of the unique library insertion types (the combination of gene number, insertion length and location). Isolates with no matches to reference either side of the hit, usually when the hit was present in a small contig, were discarded from the analysis.

Once hits had been allocated an insertion type, the node at which the insertion occurred was reconstructed on the Gubbins phylogeny for each GPSC. This ancestral state reconstruction was performed using PastML [82], as these phylogenies were not time-calibrated. The recombination predictions were then searched to detect whether there was a putative recombination event, on the branch on which acquisition was estimated to occur, spanning the insertion site within the reference for a GPSC. If there was a recombination event at this node, this was considered indicative of element insertion via homologous recombination. The flanking regions of the isolate with the fewest reconstructed SNPs around the insertion site of the element since its insertion, as inferred from the Gubbins base reconstruction, were then extracted to test for the origin of this element.

These flanking regions were compared to a reference collection of 52 streptococcal genomes collated from antimicrobial susceptible *S. pneumoniae* and other Streptococcus species, building on the database collated in Mostowy *et al* 2017 [84]. BLASTN was used to compare each flanking region to this database. The orthologous regions to these flanks were also extracted from isolates not containing the insertion, to act as a control.

The statistic *γ* was used to determine the species of origin for an insertion. This utilised the BLAST bit score, which is a normalized form of the raw score of an alignment, representing the size of the search space required to find a hit of a similar score. The *γ* statistic was calculated as the bitscore of the top ranked *S. pneumoniae* hit (b) divided by the bitscore of the top ranked hit (B):

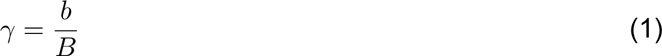

Hits where the top match was *S. pneumoniae*, indicating the insertion originated from an intraspecies transformation event, had a *γ* score of 1. Any score below 1 indicated a potential origin from outside of *S. pneumoniae*.

This pipeline was further modified to detect the origin of *pbp* genes involved in the acquisition of resistance. Here, using phenotype predictions from the RF model described above, ancestral node resistance states were reconstructed. The descendants of nodes where resistance was acquired or lost with the fewest SNPs in the three *pbp* genes then had their gene sequences extracted. These sequences were then compared to the reference database using BLASTN. The same *γ* statistic as above was used to detect the likely origin of these *pbp* genes. The code for both the altered *pbp* pipeline and the MGE detection pipeline is available at https://github.com/jdaeth274/ISA.

For *murM*, where the effect of alterations on resistance levels are less well understood, a different approach was taken. The regions corresponding to the *murM* genes in the annotated references were extracted from the PMEN3 and PMEN9 whole genome alignments. To enable the detection of possible interspecies recombinations, *murM* sequences from *Streptococcus mitis* 21/39 (accession code: AYRR01000000) and Streptococcus pseudopneumoniae IS7493 (accession code: CP002925) were added to the dataset. All *murM* sequences were then aligned with Muscle v3.8.31 [85]. Sequences were clustered into lineages, and recombinations inferred, using fastGEAR [84].

## Results

### Divergent genomic epidemiology of antibiotic-resistant pneumococci

Analysis of the PMEN3 and PMEN9 collections with Gubbins showed the lineages had distinct evolutionary and transmission histories. The phylogeny representing the evolution of PMEN3 was constructed from isolates of GPSC6, which were collected for 31 countries over 23 years (1992 to 2015; Figure 2). This range was sufficient for the estimation of a molecular clock (Figure 2-figure supplement 1). This estimated a root date of 1942 (95% credible interval of 1910 to 1959) and a molecular clock rate of 1.69 x 10^−6^ substitutions per site per year (95% credible interval of 1.52 x 10^−6^ to 1.86 x 10^−6^ substitutions per site per year). The PMEN3 phylogeny is dominated by a single 491 isolate ST156 clade, which is found in 27 countries mainly from Europe (85 isolates), North America (90 isolates) and South America (192 isolates). Most of these isolates are either of the ancestral serotype 9V, or serotype 14, with changes between these two serotypes accounting for 9 of 36 serotype switches reconstructed within the clade (Figure 2-figure supplement 2). Both of these serotypes were targeted by the PCV7 vaccine. However, there is a clade of 26 isolates of serotype 19A, not included in PCV7, from the USA with a most recent common ancestral node date of 2000 (95% credible interval of 1999 to 2001). This coincides with the date of PCV7’s introduction into the USA, consistent with these switched isolates evading the vaccine and persisting until PCV13 (which includes 19A) was introduced [49]. In total 13 serotypes are found in the collection, of which seven (11A, 13, 15A, 15B/C, 23A, 23B & 35B) are not found in the PCV13 vaccine. Hence a single PMEN3 clade has rapidly diversified its surface antigens as it has frequently disseminated between countries.

**Figure 2:**
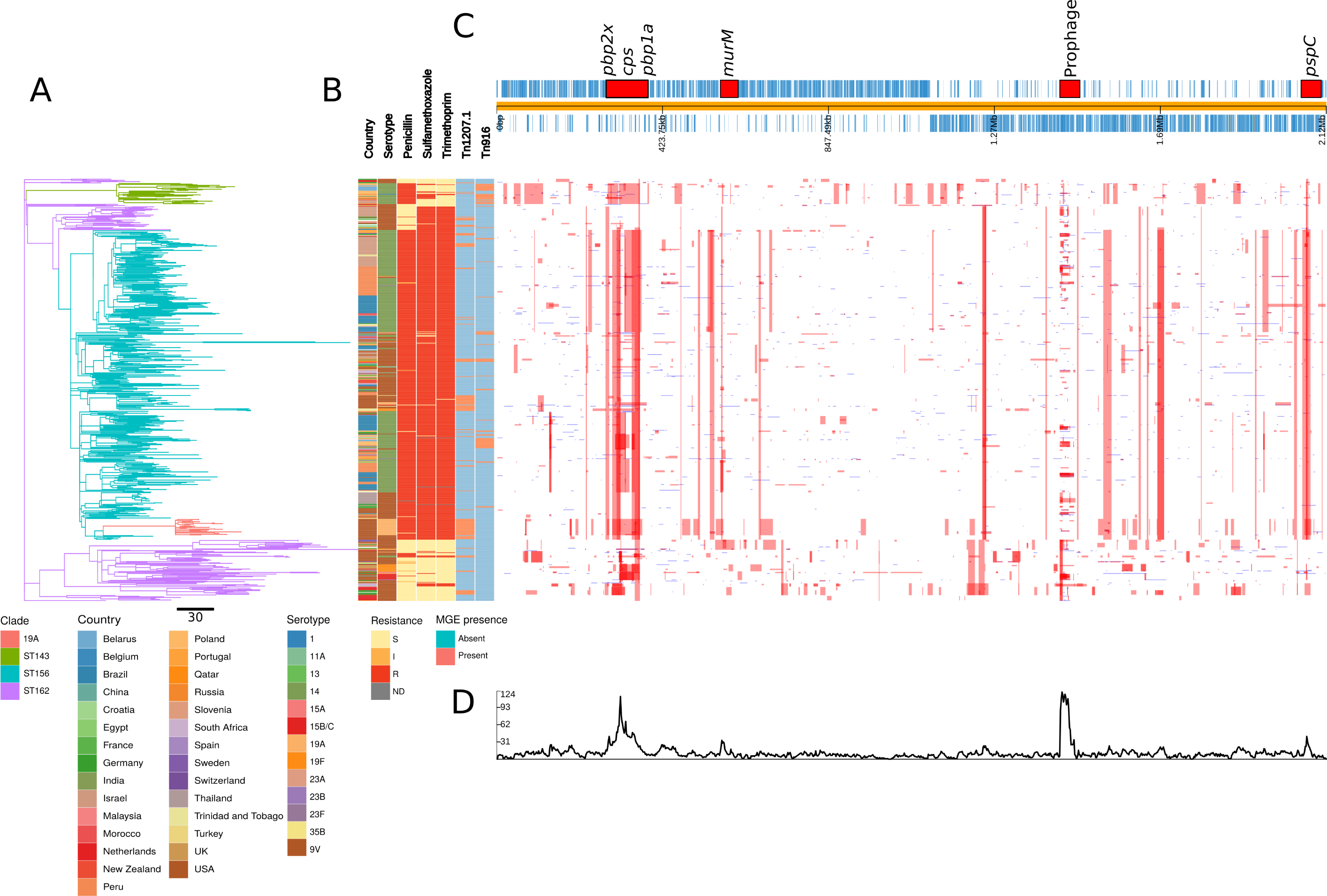
Phylogenomic analysis of PMEN3 lineage. **(A)** ML phylogeny of the non recombining regions of the PMEN3 collection. Branches are coloured by clade identified in the key. 699 isolates are present in the tree. Units for the scale bar are the number of point mutations along a branch. **(B)** Bars highlighting the country of origin, serotype, resistance categories to pencillin, trimethoprim and sulfamethoxazole and the presence of the MGEs Tn*1207.1* and Tn*916* among isolates. Bars map across to isolates on the phylogeny.**(C)** Simplified annotated genome of the PMEN3 reference isolate RMV4. Regions are highlighted are situated around peaks of recombination event frequency. Blue bars represent individual genes annotated within the assembly.**(D)** Distribution of recombination events across the PMEN3 collection. In the upper half of the graph, red bars indicate recombination events occurring on internal nodes in the tree, which are subsequently present in multiple isolates. These bars map across to isolates in the phylogeny in section *A* and map to regions in the genome annotated in section **C**. Blue bars indicate recombination events on terminal nodes of the tree, occurring in only one isolate. In the bottom half of the graph, the line represents the frequency of recombination events along the genome’s length.

By contrast, the phylogeny representing the evolution of PMEN9, constructed from isolates of GPSC18, was split into multiple clades that are separated by deep branches (Figure 3). Even when excluding the outlying serotype 7C isolates, the only discernible molecular clock signal suggested this strain was centuries old (Figure 3-figure supplement 1). Despite this age, the largest clades generally remained confined to particular countries. The largest clade was associated with Germany (accounting for 166 of the 250 isolates), with other representatives from Slovenia and China. Other clades were associated with the USA (accounting for 91 of the 98 isolates), South Africa (accounting for 68 of the of 73 isolates), and China (accounting for 18 of 45 isolates). All the isolates in the three largest clades express serotype 14, as do 93% of all isolates in this phylogeny. While in the Chinese clade, two monophyletic isolates had switched to 19F from serotype 14 and a further monophyletic pair had switched to 23F from 14. Only nine serotype switches were identified across GPSC18 (Figure 3-figure supplement 2). In total there were six serotypes present within the collection, only two of which (16F and 7C) were not found in the PCV13 vaccine. Overall, there is little evidence of frequent intercontinental transmission or antigenic diversification with this set of isolates. Hence genomics suggests different histories for these lineages, despite them both being internationally disseminated antibiotic resistant *S. pneumoniae*, commonly expressing the invasive serotype 14, with identical sampling approaches.

**Figure 3:**
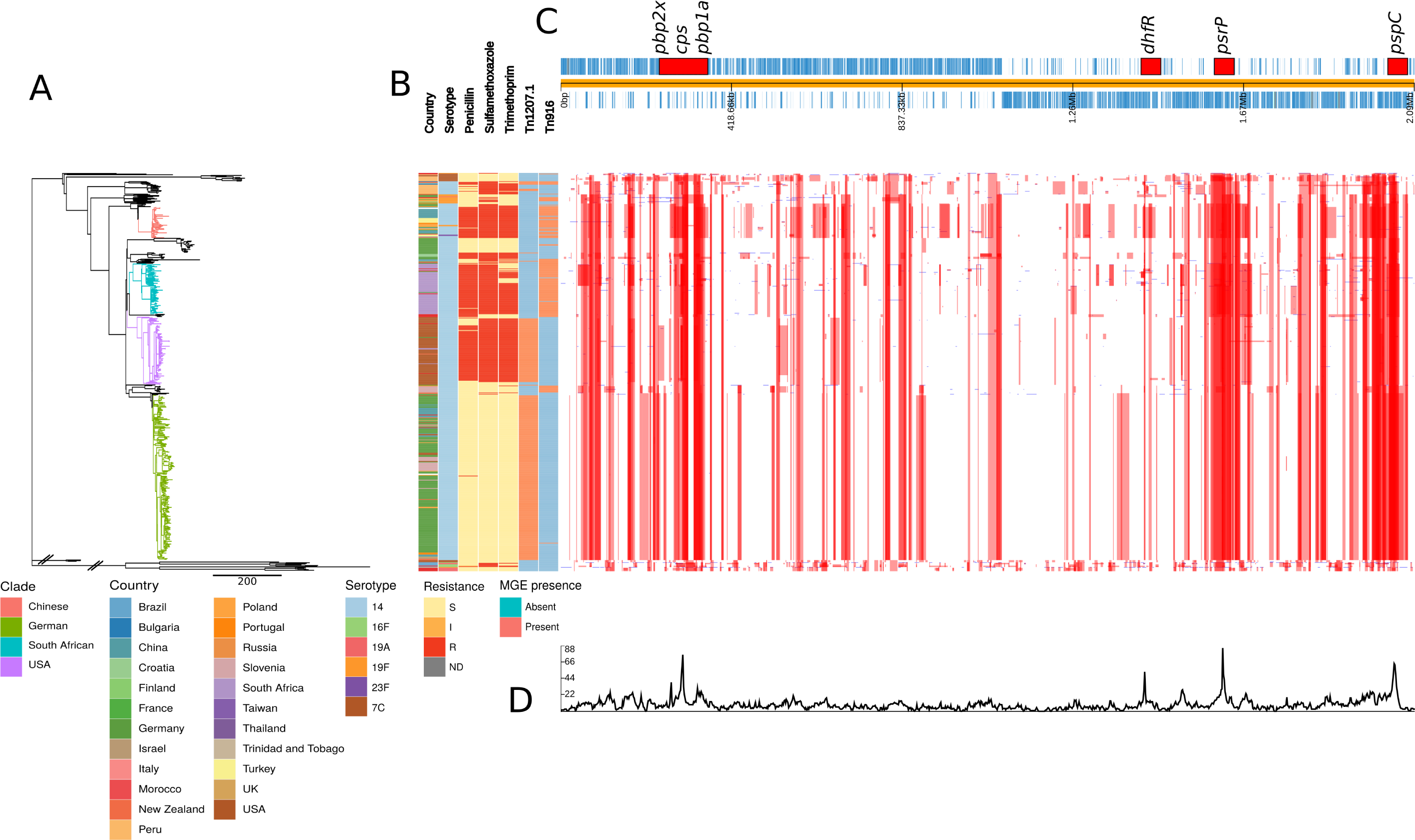
Phylogenomic analysis of PMEN9 lineage. **(A)** ML phylogeny of the non recombining regions of the PMEN9 collection. Branches are coloured by clade identified in the key. 575 isolates are present in the tree. Units for the scale bar are the number of point mutations along a branch. **(B)** Bars highlighting the country of origin, serotype, resistance categories to pencillin, trimethoprim and sulfamethoxazole and the presence of the MGEs Tn*1207.1* and Tn*916* among isolates. Bars map across to isolates on the phylogeny.**(C)** Simplified annotated genome of the PMEN3 reference isolate INV200. Regions are highlighted are situated around peaks of recombination event frequency. Blue bars represent individual genes annotated within the assembly.**(D)** Distribution of recombination events across the PMEN3 collection. In the upper half of the graph, red bars indicate recombination events occurring on internal nodes in the tree, which are subsequently present in multiple isolates. These bars map across to isolates in the phylogeny in section *A* and map to regions in the genome annotated in section **C**. Blue bars indicate recombination events on terminal nodes of the tree, occurring in only one isolate. In the bottom half of the graph, the line represents the frequency of recombination events along the genome’s length.

### Variation in transformation rates and imported sequence properties

The two lineages also differed in the patterns of recombination across their genomes. In the PMEN3 reference genome, there is a high density of recombinations around a 45 kb prophage region, indicating frequent infection by phage. Exclusion of these recombination events allowed estimation of the overall ratios of base substitutions resulting from homologous recombination relative to point mutations (r/m). Consistent with its more rapid serological diversification, r/m was higher in PMEN3 (13.1) than PMEN9 (7.7).

This difference in r/m could be due to the two lineages differing in three ways: (i) in the number of recombinations, (ii) in the length of recombination events or (iii) in the sources of their recombination events, with more divergent sources increasing the r/m.

The first explanation partially accounted for the difference: there were 0.129 recombinations per point mutation in the PMEN3 reconstruction, compared to 0.093 per points mutation in PMEN9. Comparisons of the length distribution and SNP density of recombination events revealed further differences between the two lineages (Figure 4). PMEN3 generally imported sequence with a higher SNP density, with a median SNP density of 11.8 SNP/kbp of sequence imported, compared to PMEN9, which had a median SNP density of 9.3 SNP/kbp. Therefore, the difference in r/m between the two lineages reflects both the increased frequency of recombination in the PMEN3 lineage and the increased diversity of the imported sequence.

**Figure 4:**
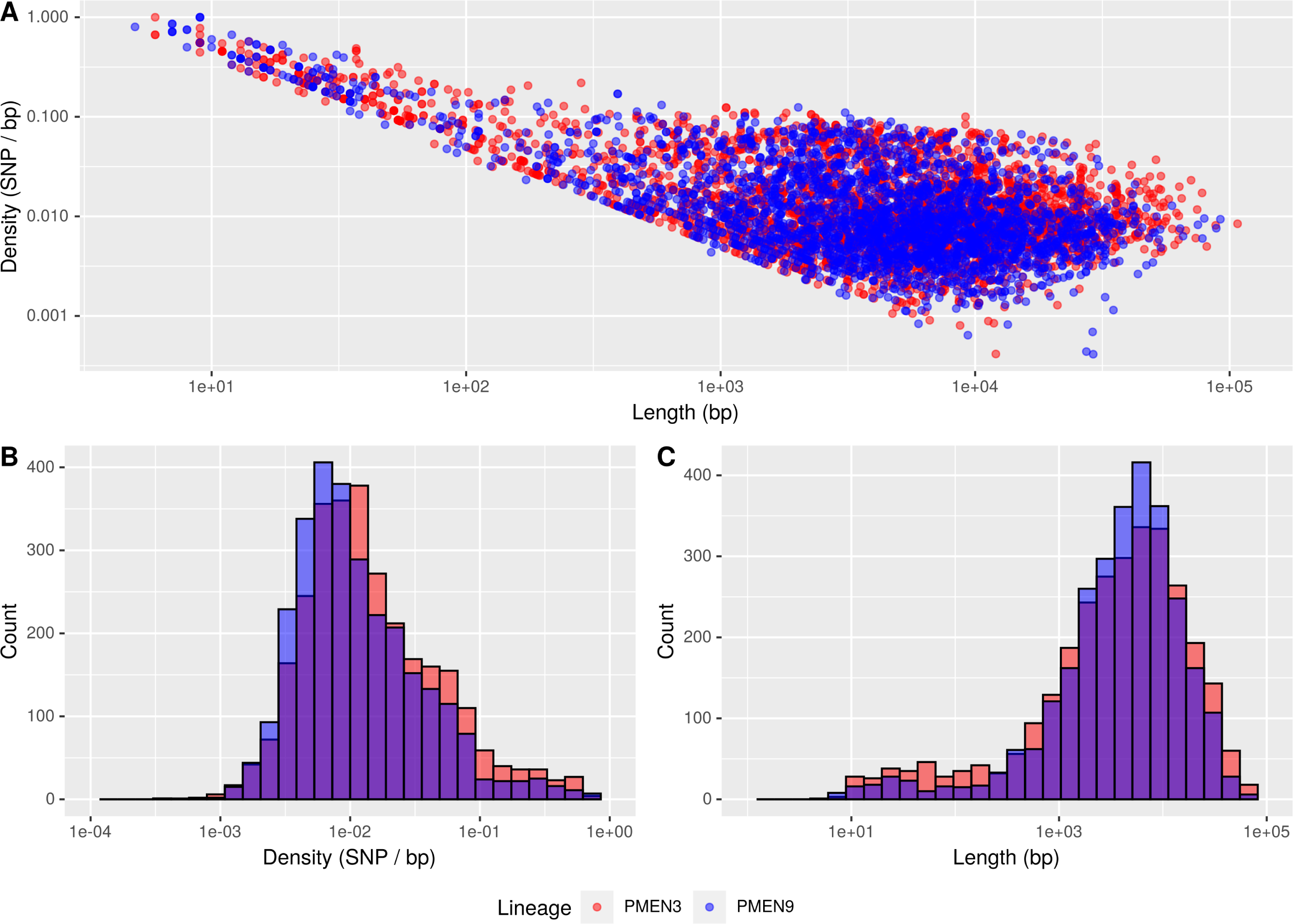
Summary of recombination differences between PMEN3 and PMEN9. A Plotting the SNP density against the Length of recombination events across the PMEN lineages. Dots represent individual recombination events and are coloured by Collection. B Overlaid histograms of the SNP density per recombination event in the PMEN collections.C Overlaid Histograms of the length of bases in each recombination event across the PMEN collections.

Several peaks of recombination within the chromosome correspond to loci likely under immune selection. In PMEN9, there is an elevated density of recombinations affecting the *psrP* gene, encoding the antigenic pneumococcal serine rich repeat surface protein. Additionally, the antigenic Pneumococcal Surface Protein C, encoded by *pspC*, is a recombination hotspot in both lineages. Both these genes are highly diverse in pneumococcal populations, and elicit strong immune responses from hosts [86].

Both strains have large recombination hotspots at their *cps* loci, which determine an isolate’s serotype. These loci undergo frequent recombination in the serologically diverse PMEN3, with fewer events at this locus in PMEN9. We tend to observe large recombination blocks around these loci, spanning the regulatory gene *wcjG* and the *wze*, *wzd*, *wzh* and *wzg* modulatory genes [31, 87] (Figures 2-figure supplement 2 & 3-figure supplement 2). Within PMEN3, for switches from 9V to 14, the median recombination block size across the 20 kb *cps* locus was 26.5 kb in length. These recombination blocks frequently encompassed the *pbp1a* and *pbp2x* genes involved in penicillin resistance. In PMEN9, three of the seven recombination events causing serotype switches spanned either *pbp1a* or *pbp2x*; this proportion increased to over 75% (26 of the 34) of the recombinations associated with a serotype switch in PMEN3.

### Emergence of beta lactam resistance through interspecies transformation

Penicillin resistance was predicted using a RF model on the PBP TPDs (see Methods), which categorized isolates using the pre-2008 CLSI meningitis resistance breakpoints (Figure 5-figure supplement 1). In the PMEN9 collection, 62% of isolates were susceptible to penicillin (recorded or predicted MIC <= 0.06 *µ*g/ml; Figure 3). However, 78% of the PMEN3 collection was classed as resistant (MIC >= 0.12 µg/ml), with 21% susceptible to penicillin, and the remaining 1% classified as intermediately resistant (0.06 µg/ml < MIC < 0.12 µg/ml; Figure 2).

Across the two PMEN lineages, there were 32 changes in resistance profile for penicillin. The most common alteration was acquisition of resistance by sensitive isolates, with 18 instances in the two collections (56% of events). There were also seven instances of resistant isolates reverting to penicillin sensitivity across the collections. In 21 of the 32 alterations in resistance profile, the evolutionary reconstruction identified at least one of the three resistance-associated *pbp* genes was altered by a concomitant recombination event.

In PMEN3, the spread of penicillin resistance was reflected by the expansion of the ST156 clade, 98% of which was penicillin resistant. Recombinations altered *pbp1a*, *pbp2b* and *pbp2x* at the base of this clade (Figure 2).

Combining the time calibrated phylogeny with the ancestral state reconstruction of penicillin resistance showed the penicillin-resistant proportion of GPSC6 increased throughout the early 1980s with the expansion of the ST156 clade, which originated in 1984 (95% credible interval 1981 to 1986) (Figure 5). This expansion of resistant lineages continues until the early 2000s, when it appears to plateau within the strain from roughly 2010 onward.

**Figure 5:**
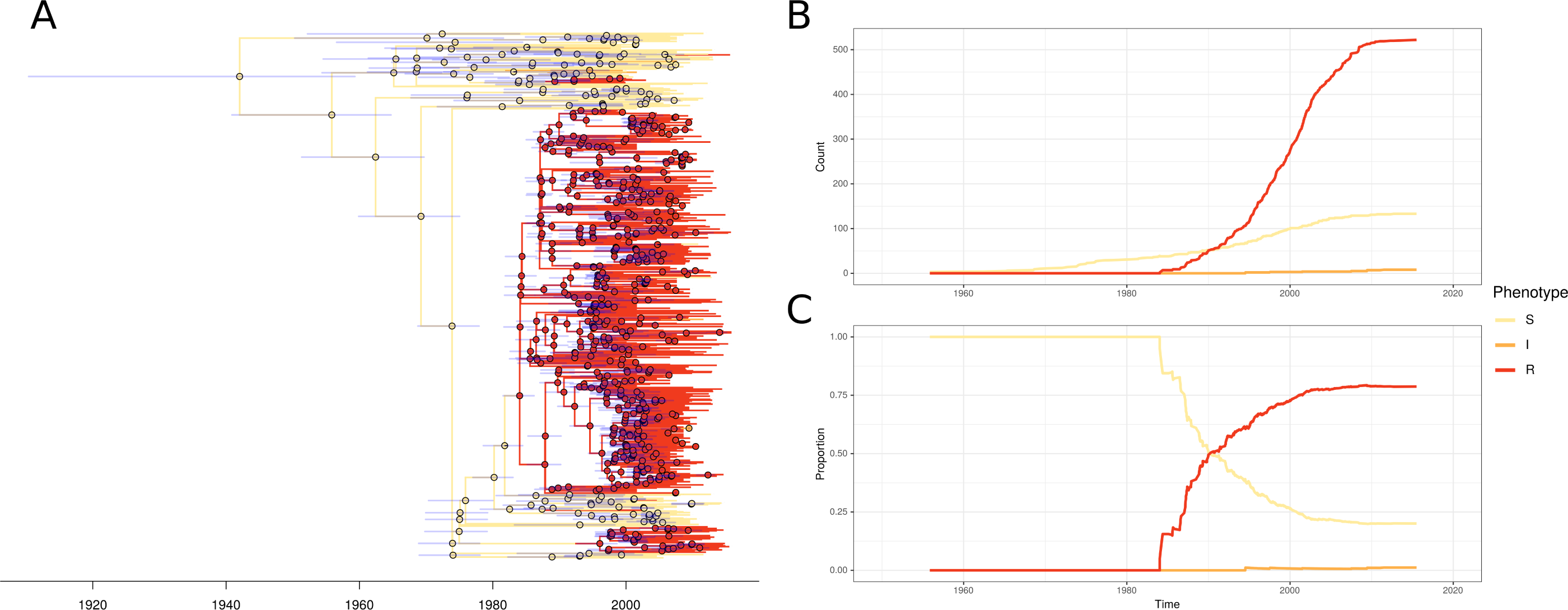
PMEN3 resistant lineages through time. **(A)**. Time calibrated phylogeny of PMEN3. Branches are coloured by inferred resistant state. Pie charts present at nodes represent the probability of each state. Blue bars across the nodes represent the 95% credible interval for the age of the node. **B** The reconstructed absolute number of branches per resistant phenotype through time. **C** The proportion of total branch over time in either of the four states.

The highest MICs within the ST156 clade (up to 8 *µ*g/ml) were associated with the vaccine-escape 19A clade of isolates from the USA (Figure 5-figure supplement 2). This reflects a 53 kb recombination spanning the cps locus, which caused the alteration in serotype, also spanning *pbp1a* and *pbp2x* (Figure 2-figure supplement 2). Hence the PCV7-escape recombination also reduced susceptibility to antibiotics. The converse situation was observed for a single ST156 clade member that had reverted to susceptibility. A 53 kb recombination event, causing a switch from serotype 9V to 15B/C (Figure 2-figure supplement 2), also restored the ancestral, susceptible versions of *pbp2x* and *pbp1a*. Conversely, in PMEN9, penicillin resistance emerged independently in different locations. The USA and South African clades appear to have gained resistance independently. There are 12 recombination events on different branches across the clade spanning *pbp2x* for these highly resistant US isolates (Figure 5-figure supplement 2). The largest of these imports 4.7 kb of sequence. Around the *pbp1a* gene there is also one large 12 kb recombination event. While for the *pbp2b* gene, there is 4.4 kb recombination spanning the length of the gene. Alteration in the *pbp2x* and *pbp2b* genes are the first steps towards resistance, although isolates with solely a mosaic *pbp2x* gene have been found to be resistant to penicillin [88].

As penicillin resistance was originally demonstrated to involve the acquisition of sequence from related commensal streptococci [34, 35, 89], the origin of these *pbp* genes within recombination events was analysed with a simple statistic, *γ* (see Methods). This had a value of one, if a recombination was likely to originate within *S. pneumoniae*, else was lower, if it came from a donor of a related species. Across the gains of resistance from penicillin sensitivity, the median *γ* score for *pbp1a* was 1.0, while for *pbp2b* it was 0.94 and for *pbp2x* it was 0.72. This pattern also applied to the emergence of ST156 (*pbp1a* = 1.0, *pbp2b* = 0.92 & *pbp2x* = 0.62), suggesting the *pbp2x* and *pbp2b* loci were most affected by recombination with non-pneumococcal streptococci. As a control, the *γ* score for the *pbp1a* and *pbp2x* genes was 1.0 for the reversion to penicillin susceptibility within ST156 (*pbp2b* was not present within a recombination block for this alteration).

Isolates from a further 146 GPSCs were also analysed for the origin of their *pbp* genes. Penicillin resistance levels across the whole GPS collection were estimated using the RF method as described for the MDR collections (see Methods). 19,956 of 20,015 (99%) isolates had successful predictions for penicillin resistance phenotype. The majority of isolates, 64%, were susceptible to penicillin, while 30% were classified as resistant and the remaining 5% intermediately resistant. Ancestral state reconstructions across these strains identified 300 alterations in penicillin resistance levels. The most common changes were susceptible to resistant, occurring 123 times (41%), and susceptible to intermediately resistant, which occurred 85 times (28%). In total, 160 of the 300 alterations (53%) were associated with an inferred recombination event affecting at least one of *pbp1a*, *pbp2b* or *pbp2x*. The *pbp2x* gene was most frequently identified as being altered by recombination, occurring in 87 of the 160 alterations associated with a recombination.

As is the case with the PMEN lineages, the emergence of resistance from susceptible genotypes was associated with at least fragments of the *pbp2x* and *pbp2b* genes often being imported from other species (indicated by *γ* < 1; Figure 5-figure supplement 3). The median *γ* score for *pbp2b* was 0.96 with *pbp2b* having a *γ* score < 1 in 23 gains of resistance. While the median *γ* score for *pbp2x* was 1, there were 15 gains of resistance where *γ* is < 1.0. The median *γ* score of *pbp1a* was 1, with only one instance where its score was < 1.0. However, where resistant isolates reverted to susceptibility, across all three genes the median *γ* score was 1.0, indicating within species recombinations could drive loss of resistance.

### Evolution of resistance through recombination at other core loci

In PMEN3, there were further peaks in recombination frequency around the *murM* gene, which encodes an enzyme involved in cell wall biosynthesis [90], and has also been implicated in mediating penicillin resistance [7, 91]. Yet compared to the *pbp* genes, the relationship between *murM* modifications and penicillin resistance is much less precisely characterised. Therefore an alignment of the *murM* sequences was analysed with fastGEAR [84] to identify any patterns of sequence import from related species that may be associated with penicillin resistance (Figure 5-figure supplement 4). This revealed evidence of recombination with *S. pseudopneumoniae* and *S. mitis* at *murM* in both lineages. However, only one modification, affecting the region 946 bp to 1143 bp within *murM*, was associated with high-level penicillin resistance. This alteration was observed in both the PMEN9 USA clade and the PMEN3 19A clade, which exhibited the highest penicillin MICs in the two datasets (Figure 5-figure supplement 2).

In PMEN9, there is a high density of recombination events affecting the *dhfR* gene, the sequence of which determines resistance to trimethoprim, one of the two components (along with sulfamethoxazle) of co-trimoxazole [92]. However, PMEN9 is largely trimethoprim and sulfamethoxazole susceptible, with 60% and 54% of isolates predicted to be susceptible respectively (Figure 3). Yet, within the South African, Chinese and USA clades there are high levels of resistance to both trimethoprim and sulfamethoxazole. Within the South African clade, 99% are resistant to sulfamethoxazole, 77% are resistant to trimethoprim and 77% resistant to cotrimoxazole, in the USA clade 94% of isolates are resistant to all three and in the Chinese clade all isolates are resistant to all three. Within South Africa, these high resistance levels could be driven by widespread co-trimoxazole use, as it is commonly used as a prophylactic treatment against secondary infections in HIV positive individuals [93].

Trimethoprim and sulfamethoxazole resistance was much more widespread among the PMEN3 collection. In total 80% of isolates within PMEN3 were trimethoprim resistant and 81% were sulfamethoxazole resistant (Figure 2). This spread was mainly driven by the expansion of the ST156 clade, which inherited alleles conferring these resistance phenotypes. By contrast, within the ST143 clade, only 12% of isolates are trimethoprim resistant and 36% are sulfamethoxazole resistant. The 15 isolates from South Africa in the collection are all resistant to both trimethoprim and sulfamethoxazole. Hence co-trimoxazole resistance spreads through clonal expansion in both lineages, albeit to a much greater extent in PMEN3.

Levels of resistance to trimethoprim and sulfamethoxazole were high across the GPS collection. A majority of isolates were resistant to sulfamethoxazole (11,576 of 20,015; 58%), with fewer isolates resistant to trimethoprim (7,765 of 20,015; 39%). The combination of resistances, conferring full cotrimoxazole resistance, was identified in 7,661 isolates (38%).

### MGE spread among MDR collections

Other antibiotic resistance phenotypes are determined by acquired genes, rather than alterations to the sequences of core genes. Two resistance associated MGEs were widespread in PMEN3 and PMEN9: Tn*916*, an ICE encoding *tetM* for tetracycline resistance; and Tn*1207.1*, a transposon encoding a *mef(A)*/*mel* efflux pump causing macrolide resistance [16, 94].

Tn*916* was present in 70 representatives of GPSC6 (PMEN3) (Figure 2). An ancestral state reconstruction identified 17 independent insertions. Only two spread to a notable extent: one was a clade of 22 isolates within the ST156 clade, and the other was ST143. However, there were multiple instances of Tn*916* being lost, by five and 13 isolates in each clade, respectively. In PMEN9, Tn*916* was present in 150 isolates (Figure 3). The most common was in the South African clade, where Tn*916* was found in 71 of the 73 isolates in this clade, with likely deletion in two isolates. Similarly, 40 of the 45 isolates within the Chinese clade had also acquired Tn*916*, with five isolates without the element appearing to have lost Tn*916* independently. There was also a smaller insertion in 10 isolates outside the German clade, and then further sporadic insertions of Tn*916* around the phylogeny.

Tn*1207.1* was more common in both strains. It was found in 108 isolates of PMEN3 (Figure 2), resulting from 27 independent insertions. The two insertions associated with the largest clonal expansions were one within the 19A subclade (26 isolates), and a second in another subclade of the ST156 clade (25 isolates). The other 25 insertions were less successful, appearing sporadically around the phylogeny. In PMEN9, Tn*1207.1* was present in 341 isolates (Figure 3). The element was present in 92 isolates of a subclade of the USA clade, and ubiquitous in the 238 isolates of the German clade, the most successful insertion observed in the collection Hence both elements were acquired on multiple occasions by both lineages, suggesting frequent importation. However, few of these insertions resulted in internationally disseminated antibiotic resistant pneumococcal genotypes.

### Selection for local expansion of macrolide resistant *S. pneumoniae*

The expansion of the German clade carrying Tn*1207.1* represented an unusual case of an MGE insertion being associated with a successful genotype. This suggested strong selection for a macrolide resistant genotype in Germany in recent years. However, German antibiotic consumption is generally low relative to the rest of Europe. Notably, the German PMEN9 clade is penicillin susceptible. Based on macrolide and penicillin consumption data for the period from 1992 to 2010, Germany has a high ratio of macrolide to penicillin usage relative to other European countries (Figure 6-figure supplement 1).

A phylodynamic analysis of the 162 German isolates in this clade was used to test whether this atypical pattern of antibiotic consumption could explain the success of the clade. There was significant evidence of a molecular clock, based on the correlation between the root to tip distance and the date of isolation for this clade (n = 162, R^2^ = 0.15, p value < 1 x 10*^−^*^4^) (Figure 6-figure supplement 2) . This estimated the clade’s most recent common ancestor (MRCA) existed in 1970. Generating a time calibrated phylogeny using BactDating suggested a relatively slow clock rate of 5.5 x 10*^−^*^7^ substitutions per site per year (95% credibility interval of 4.5 x 10*^−^*^7^ to 6.6 x 10*^−^*^7^ substitutions per site per year).

The Skygrowth package was then used to reconstruct the effective population size, N*_e_*, and the growth rate of N*_e_* through time of this clade. The antibiotic usage data over this period was used as a covariate, to test for evidence of selection by changing consumption (Figure 6). All these isolates are serotype 14, which is included in the PCV7 vaccine that was introduced into the universal vaccination programme for children under two years of age in Germany in 2006 [95]. As such, only isolates collected pre-2006 were included in this analysis, to minimise any effect of the vaccine on N*_e_*.

**Figure 6:**
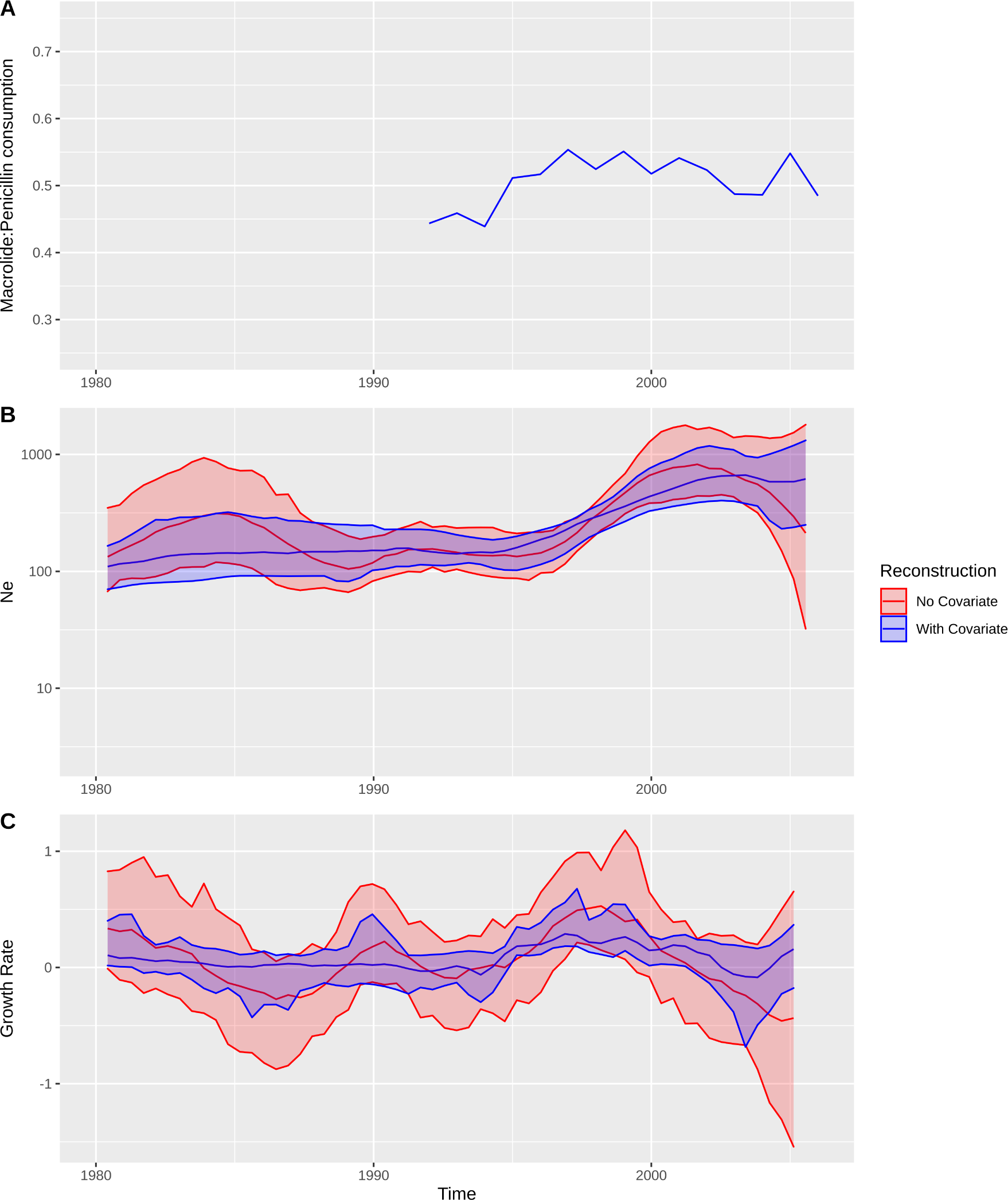
Expansion of a macrolide resistant clade in Germany pre-vaccine. **(A)** The ratio of macrolide to penicillin consumption in Germany. **B** The change in N*_e_* through time inferred by skygrowth, with the red line representative of when no covariates are incorporated and the blue line when macrolide use is incorporated into the reconstruction. Shaded regions represent the 95% credible intervals. **C** The reconstruction of the growth rate of N*_e_* through time. The red line representative of when no covariates are incorporated, and the blue line when macrolide consumption data are incorporated. Shaded regions represent the 95% credible interval for the reconstruction.

From the reconstruction without using the macrolide to penicillin data, we can see that this lineage expanded rapidly during the late 1990s and early 2000s, with its peak in growth rate around 1997 preceding a peak in N*_e_* around 2003. Following this peak there is a decline in both N*_e_* and growth rate.

When incorporating macrolide consumption data into the reconstruction, we do observe a narrowing of the credible intervals for the growth rate estimation, which is indicative of the macrolide to penicillin usage being informative. Indeed, macrolide usage has a significant positive mean posterior effect of +0.21 [95% confidence interval 0.015 to 0.50] on the growth rate of the clade [75]. This supports the hypothesis that growth rate is correlated with the consumption of antibiotics, which is consistent with selection pressures from national-level prescribing practices driving the expansion of this clade. The peak of the macrolide to penicillin consumption ratio is in the mid-to-late 1990s, whereas the peak N*_e_* is not reached until after 2000. Yet the growth rate of N*e* peaks around when the consumption ratio is highest, consistent with observed trends in *Staphylococcus aureus*, suggesting the antibiotic use pattern drove an expansion of this clade in the late 1990s [75].

The absence of penicillin resistance, or vaccine evasion through serotype switching [96], may be a consequence of the Tn*1207.1* element itself. This MGE inserted into, and split, the gene *comEC* (Figure 6-figure supplement 3), which encodes a membrane channel protein integral to extracellular DNA uptake during competence [97]. Therefore these cells were unable to exchange DNA via transformation, necessary for serotype switching and the acquisition of penicillin resistance alleles of the *pbp* genes [19]. The impact of this insertion is evident from the absence of ongoing transformation within the German clade (Figure 3).

Analysis of the origin of this MGE, via analysing the flanking regions as for the *pbp* genes above, revealed a probable interspecies origin. The flanking regions immediately adjacent to the insertion have a low percent identity matched to other pneumococci of between 92% and 94% (Figure 6-figure supplement 3). The immediate upstream 500 bp region most closely matched to a *S. mitis* reference genome (accession code AFQV00000000). Therefore other acquisitions of the common MGEs, Tn*1207.1* and Tn*916*, were analysed to determine whether they had also been recently imported from related commensal species.

### Multiple independent acquisitions of resistance genes in *S. pneumoniae*

We first identified the set of Tn*916* and Tn*1207.1* insertion sites in *S. pneumoniae*, using the 20,015 genomes from the Global Pneumococcal Sequencing project. The genomes were searched for these two elements, and hits were categorised into unique insertion types to identify the genomic locus at which they integrated. The branch of the phylogeny on which these insertions occurred was then identified, allowing the determination of whether an element was gained via homologous recombination (see Methods).

At least one of the elements was found in 5,796 isolates (29%) across 146 different GPSCs. Of these, 1,300 isolates contained both Tn*1207.1* and Tn*916* (6%). The Tn*1207.1* element was found across 64 GPSCs in 1,935 isolates (10%). The mean prevalence of Tn*1207.1* in GPSCs in which it was present was 16%. Of the 1,935 isolates containing the element, 1,743 (90%) had their insertion point reconstructed. The majority of the 192 isolates where the insertion point was not reconstructed had the Tn*1207.1* element present within a small contig with no flanking hits to the reference (104 of 197). There were 32 unique reconstructed insertion types (a specific combination of MGE length and insertion location) of the Tn*1207.1* element, spanning 22 different insertion loci (Figure 7). Some insertion loci were targeted by multiple insertion types. The loci surrounding the *rlmCD* gene, encoding a methyltransferase, was the most common target, with 7 different Tn*1207.1* insertion types targeting this region. The most common insertion type was within a Tn*916*-like element, which occurred in 1,011 (58%) of the isolates. Hence the diversity of Tn*1207.1* insertion types was relatively low, with a Simpson’s diversity index of 0.63.

**Figure 7:**
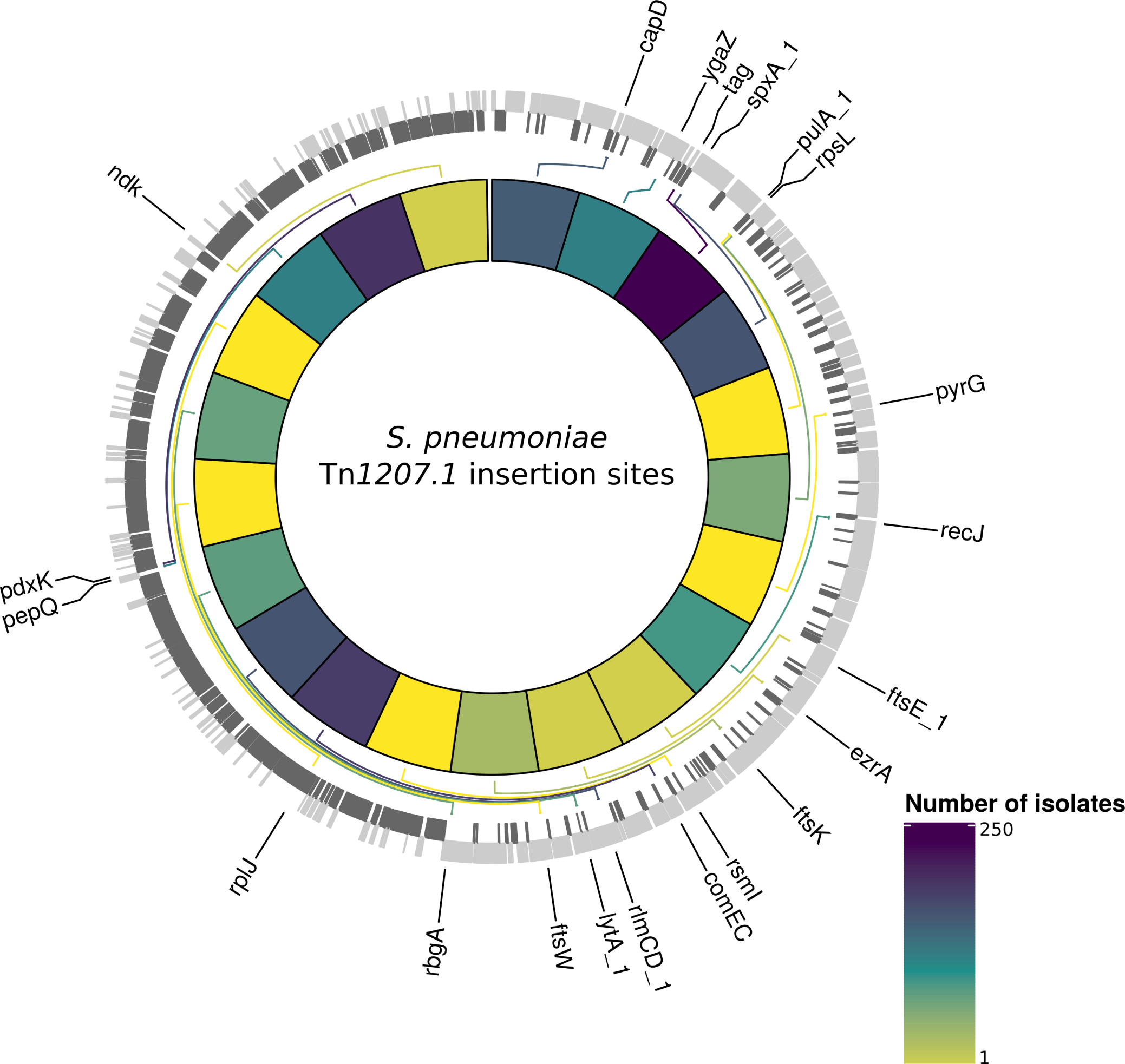
Insertion points of classified Tn*1207.1* hits within *S. pneumoniae.* Annotated genome of the reference *Streptococcus pneumoniae* (ENA accession number: ERS1681526) with genes where Tn*1207.1* has inserted either within or adjacent to among the collection. Only genes present within this element free reference are annotated. Grey bars represent coding sequences (CDS), lighter grey regions are CDS annotated in the forward strand, darker grey in the reverse. The inner heat map represents the number of isolates that have hits inserted within or adjacent to the annotated genes. The scale is log transformed.

The next most common insertion type for Tn*1207.1* was the 5.5 kb Mega version of the element inserting into, and splitting, *tag* [98]. The *tag* gene encodes a methyladenine glycosylase, involved in DNA base excision repair. This was present in 261 isolates (15% of identified hits) across 30 different GPSCs. This was also common in the PMEN collections, with Tn*1207.1* within the USA clade of PMEN9 being in the form of Mega splitting *tag* (Figure 7-figure supplement 1). The insertion of the 7.2 kb Tn*1207.1* element into *comEC*, as in the German PMEN9 clade, was the third most common, accounting for 5% of insertions in the GPS collection and appearing in 4 different GPSCs.

Contrary to the results for PMEN3 and PMEN9, Tn*916* was more widespread than Tn*1207.1* among the collection, present in 5,167 isolates across 134 of the 146 GPSCs. The mean prevalence for Tn*916* was 41% among GPSCs where it was present. Of these hits, 2,806 (54%) were not classifiable. This was primarily due to elements being present in contigs with no or very small matches back to the reference (1,239 isolates) and elements within clusters where all isolates contain the element (1,072), meaning there are no descendents of the genotype lacking the element, thereby preventing us inferring any recombination by which it may have been imported. For the classifiable 2,361 (46%) isolates, there were a large number of reconstructed insertion types, with 222 unique library hits encompassing single Tn*916* element insertions and the wider Tn*916* family of elements (Figure 8). This gives Tn*916* insertion types a Simpson’s diversity index of 0.94, indicating Tn*916* is much more variable in how it integrates into the *S. pneumoniae* genome.

**Figure 8:**
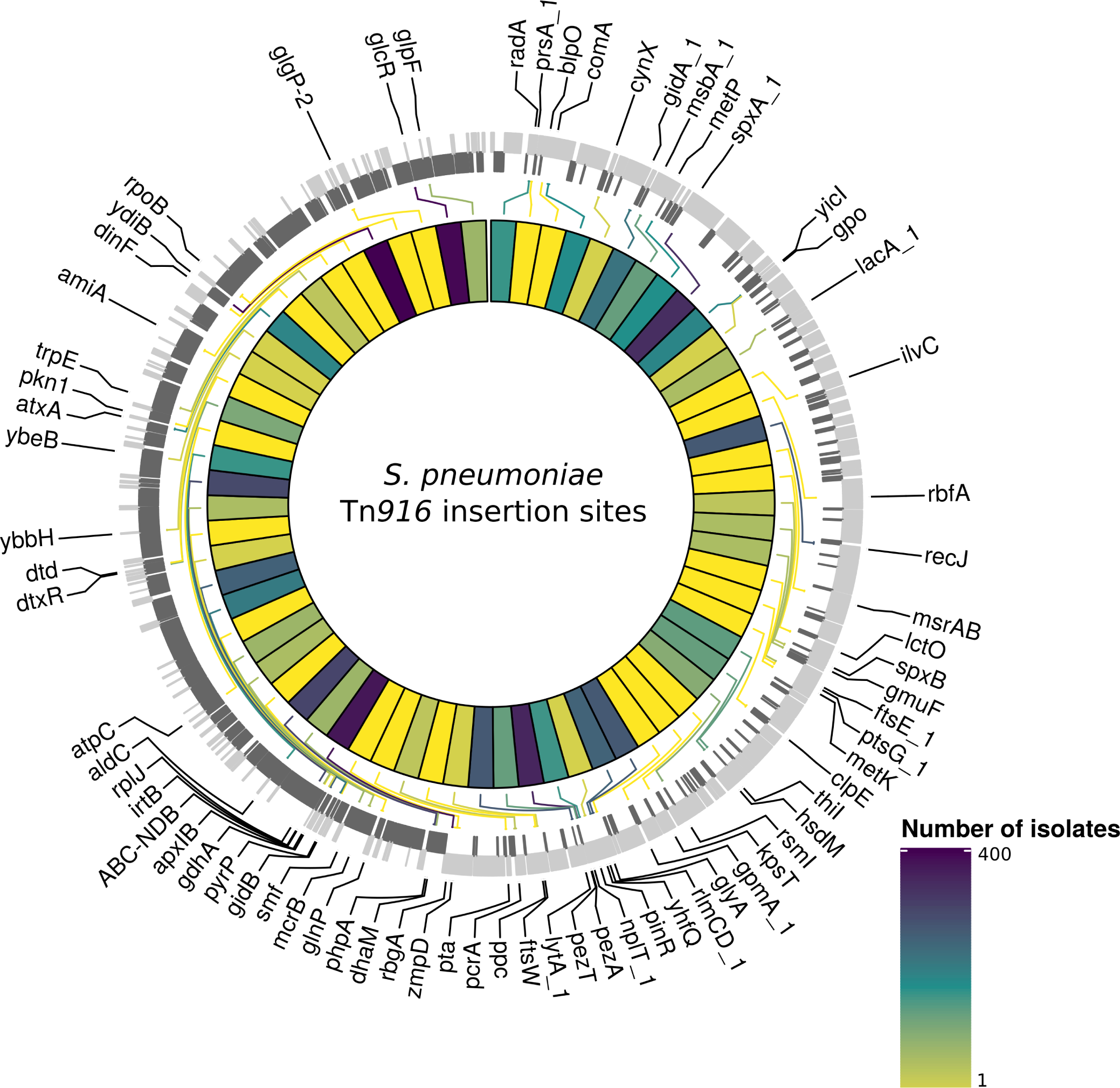
Insertion points of classified Tn*916* hits within *S. pneumoniae.* Annotated genome of the reference *Streptococcus pneumoniae* (ENA accession number: ERS1681526) with genes where Tn*916* has inserted either within or adjacent to among the collection. Grey bars represent coding sequences (CDS), lighter grey regions are CDS annotated in the forward strand, darker grey in the reverse. The inner heat map represents the number of isolates that have hits inserted within or adjacent to each of the annotated genes. The scale is log transformed.

The joint top hits for Tn*916* were insertions of the Tn*916*-like Tn*2010* and Tn*2009* elements that also contain the Mega form of Tn*1207.1*. Both insertions are present in 356 isolates each, with the majority of these occurring within the very common GPSC1 (PMEN14): 354 of Tn*2010* and 351 of Tn*2009* [40, 41]. Other common Tn*916* insertion types included a Tn*6002* element, and a much larger 84 kb element containing Tn*2009*. Tn*6002* consists of a Tn*916* backbone with an *ermB* cassette between orf20 and orf19. Tn*6002* is present in 107 isolates within a single strain. The large 84 kb insertion is a composite of a Tn*2009* and a Ωcat element. This element thus confers tetracycline, macrolide and chloramphenicol resistance, and is present in 59 isolates, also solely within one strain. Tn*916* is present in the 64.5 kb Tn*5253* in eight isolates in the collection across four different GPSCs. In general Tn*916* is present in elements over 50 kb in length in 660 isolates (28% of classifiable hits).

### Diverse insertion sites of MGEs

Ancestral state reconstruction was used to identify the insertions of Tn*916* and Tn*1207.1* across the GPS collection. For Tn*1207.1* there were 183 independent insertions of the element in its 32 identified hit types, across 55 GPSCs. The most frequent insertion type was the shorter 4.5 kb element splitting the *tag* gene, which was found to have inserted 73 times (40% of all insertions).

For Tn*916* there were a much larger number of insertion events: 585 across 93 different GPSCs. Several insertion types appeared multiple times across the collection. The most frequent (21 times) was a 72 kb element, containing only *tetM* as a resistance gene inserted upstream of *rlmCD*. The second most frequent was a 50 kb element, also only containing *tetM* as an resistance gene. This inserted 20 times around the collection, downstream of *zmpA* which encodes an immunoglobulin protease [99].

The numbers of insertions within putative recombinations differed between the two elements. For Tn*1207.1*, 64% of insertions were within recombination blocks (118 of 183), compared with only 9% of the insertions for Tn*916* (53 of 585). This difference could be driven by a couple of factors. Tn*916* encodes for its own conjugative machinery, and is often present within larger conjugative elements, and therefore may move independently of transformation. Alternatively, Tn*916* may be imported through transformation, but then transpose between loci once in a cell, thus moving away from its site of insertion. The median Simpson’s diversity index for within-GPSC Tn*916* insertion site diversity was 0.41, whereas for Tn*1207.1* it was 0.02. This suggests Tn*916*, once inserted, is likely to be able to excise and transpose within the chromosome at higher rate than Tn*1207.1*.

Recombinations mediated by transformation are generally much shorter than the lengths of these elements (Figure 9), and such exchanges generally favour deletion of elements rather than insertion [37].

**Figure 9:**
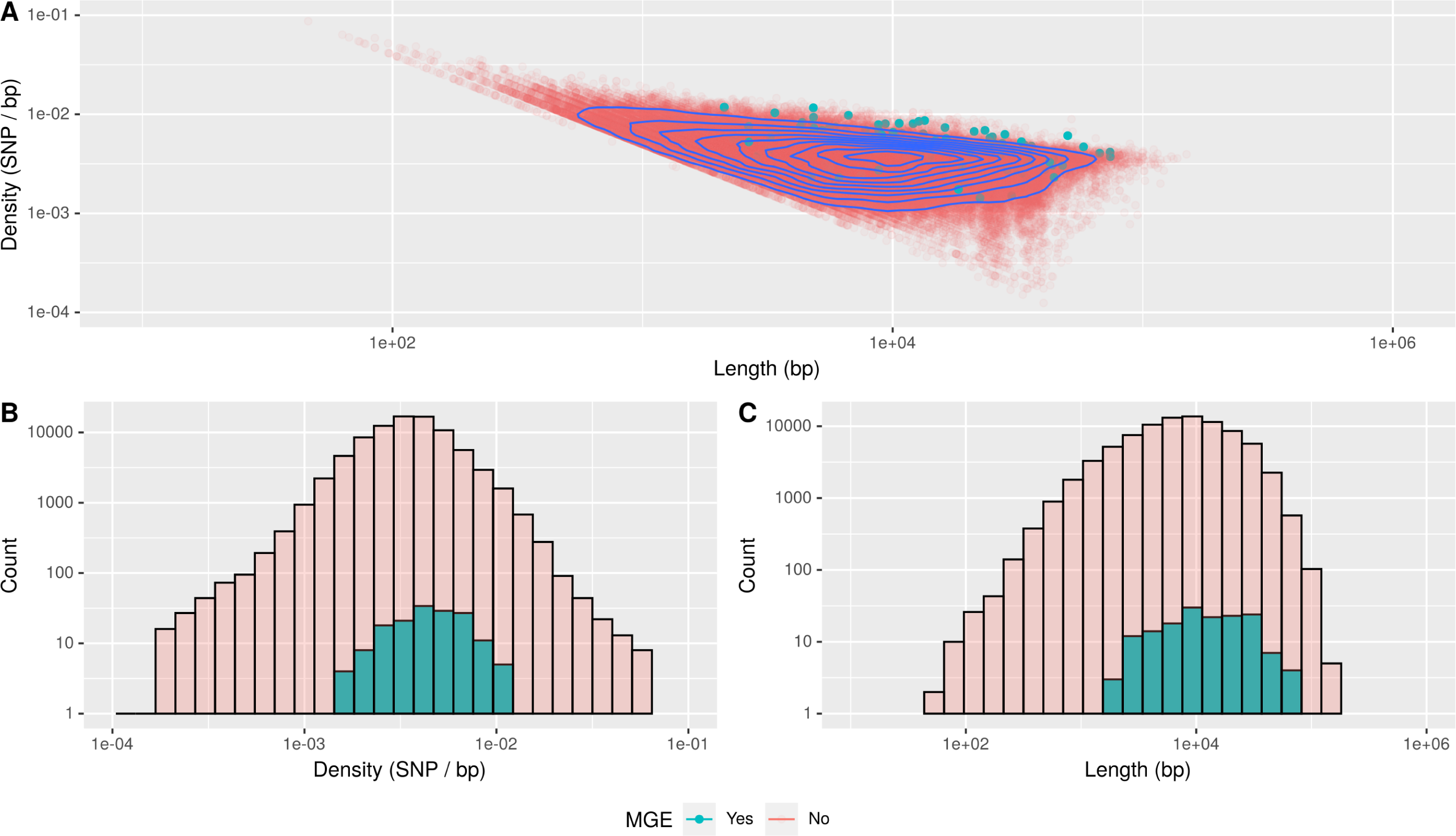
Comparison of length and SNP density of recombination events. **(A)** Joint plot of the SNP density and number of bases within recombination events for non-MGE importation and MGE importation. Blue contour lines represent the density of points present. **B** Overlaid histogram of SNP density within recombination events comparison. **C** Overlaid hisogram of the length of recombination events in bases.

Comparisons of the length distribution and SNP density for recombination events that import one of the MGEs, against other recombination events, suggested they were atypical (Figure 9). MGE recombinations were significantly longer, with a median length of 11,797 bp, compared to a median length of 7,499 bp for non-MGE recombinations and a median difference between MGE and non-MGE recombinations of 3,828 bp (Mann-Whitney U test; U = 8634645, n1 = 157, n2 = 85262, p = 3.178 x 10*^−^*^10^, 95% confidence interval 2600 to 5183 bp). Additionally, the median SNP density was significantly higher for MGE recombinations, at 4.54 SNPs per kb, compared to non-MGE recombinations, with a median of 3.53 SNPs per kb and a median difference between the densities of 0.94 SNPs per kb (Mann-Whitney U test; U =8631593, n1 = 157, n2 = 85262, p = 3.387 x 10*^−^*^10^, 95% confidence interval 0.66 to 1.27 SNPs per kb).

Given the pneumococcus tends to be conserved at core genome loci, the higher SNP density of these transformation events inserting MGEs suggested they may arise from donors of other species [61].

### Interspecies origin of MGEs

Correspondingly, analysis of the flanking regions of MGE inserts reveals a large number of hits mapping more closely to non-pneumococcal streptococci. For Tn*1207.1*, the median *γ* score was 0.89 for insertions across the flanking lengths and insertion types. For control isolates, where the element was not inserted and the orthologous flanking regions were extracted, the median *γ* score was 1.0. Hence this lower score for MGE flanks compared to homologous regions in isolates without the MGE, likely represents MGEs being acquired from other species. A Mann-Whitney U test also revealed significant difference between the control and MGE isolates *γ* scores for Tn*1207.1* (U = 2501856, n_1_ = n_2_ = 3540, p < 2.2 x 10*^−^*^16^), with the median difference between the MGE flanks score and the orthologous flanks score being -0.09 (95% confidence interval -0.1 to -0.08).

For Tn*916* insertions within recombination blocks, the median *γ* scores for both control and MGE isolates was 1. However, a Mann-Whitney U test revealed significant difference between the control and MGE isolates *γ* scores (U = 785412, n_1_ = 1524, n_2_ = 1520, p < 2.2 x 10*^−^*^16^), with the median difference between the MGE flanks score and the orthologous flanks score being -5.6 x 10*^−^*^5^ (95% confidence interval -3.4 x 10*^−^*^5^ to -5.0 x 10*^−^*^6^).

The trends in top species match for flanking regions, over increasing flanking length, follow expectations for interspecies transfers (Figure 10). The control flanking regions matched most closely to pneumococci at all tested lengths. For MGE insertion flanks, non-pneumococcal species matches were much more frequent closer to the insertion. As the flank length increased from 500 bp to 7500 bp, and linkage to the integrated resistance gene decreases, the sequence more frequently matched pneumococcal DNA.

**Figure 10:**
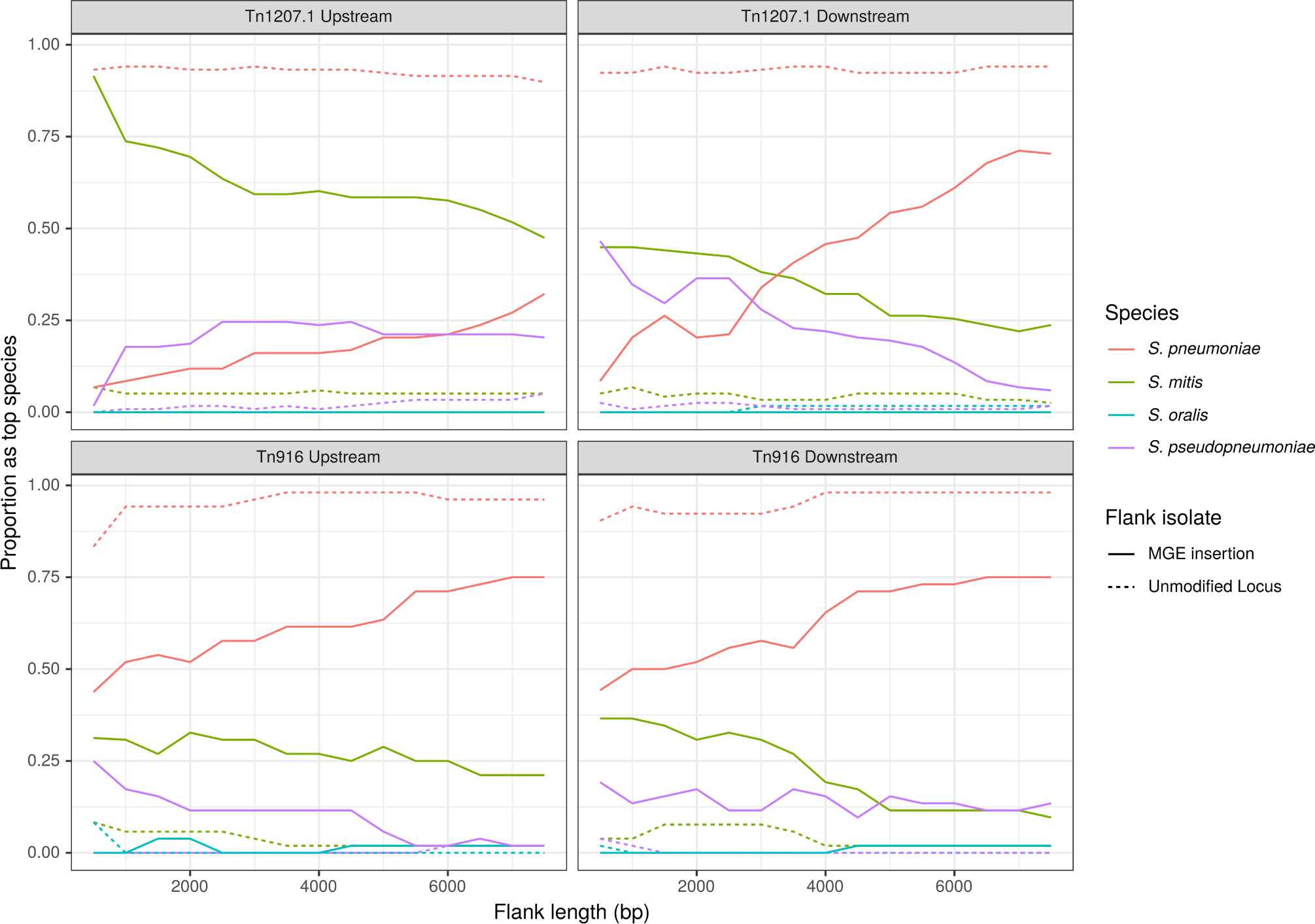
The top matching species to flanks extracted around an MGE insertion. Flanking regions upstream and downstream from MGE insertions sites were compared to a reference streptococcal database. Lines represent the proportion of matches, across all inserts reconstructed to have occurred in recombinations, that correspond to the four species present in the reference streptococcal database. The homologous regions in isolates without the MGE insertion have been extracted for comparison to these MGE hits, these are represented by the dashed unmodified locus lines. These proportions are calcuated over increasing flank length away from the inseriton site.

For Tn*1207.1*, it appeared *S. mitis* is the most likely donor, based on the results at the shortest flanking lengths (Figure 10). In the regions upstream of the Tn*1207.1* insertion *S. mitis* was the top match for 92% of 500 bp long flanks. Even at longer flank lengths *S. mitis* was still the leading match for upstream regions, although for downstream region the pneumococcus tended to become the predominant match to flanks by 4000 bp outside of the insertion.

The most common Tn*1207.1* insertion, that splitting the *tag* gene, can be used to illustrate the local import of sequence from another species (Figure 10-figure supplement 1). For the downstream flanks (Figure 10-figure supplement 1B) this trend appeared roughly linear with increasing flank length, the evidence for imported *S. mitis* sequence lost after 4000 bp. However, for the upstream flanking regions (Figure 10-figure supplement 1A), the median *γ* score remained low with increasing flank length, with a median of 0.84 at 7500 bp upstream of the insertion. This upstream region, replaced by *S. mitis* sequence in many isolates, leads into the *uvrA* gene, another component of the nucleotide excision repair machinery within the pneumococcus. The consistency of top matches for this *tag* Tn*1207.1* insertion type was high. For the upstream flank region at 500 bp long, 86% of the insertions (57 of the 66 within recombination blocks) had the *S. mitis* 21/39 (accession code AYRR00000000) reference as their top hit. In total 97% of these insertions (64 of 66) had their top hit as an *S. mitis* sequence. This lack of diversity suggests these imports originated from a single insertion in the *S. mitis* population. These imports are then likely to have moved between pneumococci multiple times.

The signal for the interspecies origins of Tn*916* insertions was less pronounced (Figure 10). While the pneumococcus tends to be the most frequent match for the regions flanking Tn*916* insertions, the proportion of matches is still far lower than seen in the control isolates, suggesting a detectable contribution of interspecies import. The difficulty of identifying insertion sites for this larger MGE suggested some interspecies transfers may have been missed. Therefore a number of Tn*916* acquisitions by the PMEN3 lineage were analysed manually. This found evidence for homologous recombination importing the elements at multiple sites, with highly divergent flanking regions at each position suggesting import from another species. This applied to independent insertions near *recJ* (Figure 8-figure supplement 1), *gmuF*, which encodes mannose-6-phosphate isomerase (also known as *manA*; Figure 8-figure supplement 2), and *gidB* (Figure 8-figure supplement 3). Additionally, Tn*916* was acquired through homologous recombination near *rplL*, with flanking remnants of Tn*5252*, on three independent occasions within PMEN3 (Figures 8-figure supplement 4 8-figure supplement 6). This suggest that acquisition of Tn*916* through interspecies transformation is sufficiently important as to detectably occur on multiple occasions within a single strain.

## Discussion

These analyses describe the evolutionary histories of the *S. pneumoniae* PMEN3 and PMEN9 lineages. Comparisons between the pair illustrate the variable epidemiology of common antibiotic resistant pneumococci. In PMEN3, the penicillin resistant ST156 clade emerged in the early 1980s, and rapidly spread worldwide. This resembles the rapid spread of PMEN1 and PMEN14 [24, 40]. In PMEN9, resistance emerged multiple times, but in clades that remained geographically confined to Germany, South Africa and the USA. This is despite the GPSC18 strain, from which PMEN9 emerged, originating earlier than PMEN3. While both lineages contain examples of acquisitions of resistance leading to successful clade expansions, there are many unsuccessful acquisitions too. The sporadic distribution of Tn*1207.1* and Tn*916* across both lineages suggests any barrier to their acquisition is not the rate-limiting factor in their spread, but rather their prevalence is limited by them being insufficiently beneficial to outcompete isolates lacking these resistances [100, 101]. This is also consistent with the intermittent deletion of the Tn*916* element across both lineages, and the reversion to penicillin susceptibility within PMEN3, suggesting there can be selection for loss of resistance. The results from these lineages are consistent with the observations from the GPS collection: hundreds of independent resistance acquisitions were observed across the species, but the majority of isolates remained macrolide and tetracycline sensitive. Taken together, this points to the ecology and context within which an isolate gains resistance being key determinants of successful spread.

The limited geographic range of clades within PMEN9 indicates the importance of local differences in selection pressure in determining the spread of antibiotic resistant variants. The South African and USA clades are both penicillin and co-trimoxazole resistant, whereas the German clade is sensitive to both, but resistant to macrolides. During the period of this clade’s expansion, Germany was amongst the lowest consumers of antibiotics in Europe [102]. It did however have a relatively high consumption of macrolides, especially relative to penicillin consumption (Figure 6-figure supplement 1). These seem to have been ideal conditions for a macrolide resistant, but penicillin sensitive, lineage to expand. Indeed, the results from our analysis imply a significant effect of the ratio of macrolide to penicillin usage on the increase in the growth rate of N*_e_* for this clade, with the greatest growth correlating with the steep increase in this ratio during the late 1990s. The highly invasive nature of this strain, at least when associated with serotype 14, means the rates of macrolide-resistant invasive pneumococcal disease were likely disproportionately high compared to the prevalence of the resistance phenotype in carriage [41, 103]. Such locally-confined transmission was also observed for the PMEN2 pneumococcal lineage [10], which also lacked the ability to undergo transformation, due to a gene needed for the competence system being disrupted by the insertion of an MGE.

In both cases, this condemned the clades to elimination by vaccine induced immunity, as the loss of transformability prevented vaccine evasion through serotype switching. Yet even in transformable PMEN9 clades, we observe far fewer serotype switching events than in the PMEN3 lineage. Hence the PCV7 vaccine has eliminated the PMEN9 lineage in the USA, where it was previously the most common antibiotic resistant lineage causing invasive disease [49]. By contrast, PMEN3 evaded the PCV7 vaccine through its acquisition of a serotype 19A capsule via a recombination that also elevated its penicillin MIC [104]. Following the introduction of PCV13 in the USA, the PMEN3 lineage is now mainly observed expressing serotype 35B [105].

In the German clade, the Tn*1207.1* that disrupted *comEC* was itself inserted through homologous recombination. Such an integration location is advantageous for a selfish element, as it prevents the Tn*1207.1* being removed or replaced in the German clade, while still permitting it to be acquired from these isolates by competent recipients in which it is currently absent [106]. This effectively reverses the normal bias towards deletion [29, 37], rather than insertion, of loci by transformation through a process akin to meiotic drive in eukaryotes [107].

Analysis of the regions flanking the Tn*1207.1* insertion into *comEC* indicate the element was acquired from a non-pneumococcal donor species, likely *S. mitis*. This highlights one of the key commonalities between both the antibiotic resistant clades of PMEN3 and PMEN9, in that they emerged following acquisition of resistant loci from related species through transformation. Interspecies transformation between viridans group streptococci species is known to be crucial where resistance emerges through modification of a core housekeeping gene via homologous recombination. The first detected penicillin resistant pneumococcus had mosaic *pbp* genes that originated from commensal viridians species, such as *S. mitis* and *S. oralis* [19, 34, 35, 89]. Analyses of penicillin resistant isolates from PMEN3, PMEN9 and the broader GPS collection, identified many gains of penicillin resistance associated with *pbp* genes that originated from non-pneumococcal species. This was particularly the case for *pbp2b* and *pbp2x*, which are usually the first alterations required for penicillin resistance to emerge [7]. However, substantial alterations in *pbp1a* are associated with higher levels of resistance, greater than the 0.12 *µ*g/ml used as the upper limit for the RF model [108]. Hence the lack of strong evidence for *pbp1a* being modified by sequence from other species may be an artefact of how transitions between discrete resistance levels were identified in this study. This may also apply to the alterations to *murM*. Particular imports from other streptococci were found in PMEN3 and PMEN9 clades with high-level penicillin resistance, which also had modified *pbp* genes, suggesting epistatic interactions are likely to be important in fully understanding the role of these imported segments [109].

These mosaic *pbp* and *murM* gene structures likely arose from short imports favoured by transformation, then integrated via homologous recombination [110]. Outside of the *pbp* loci, exchange of short DNA fragments between streptococci have been seen across the genome, especially at the competence loci [111]. Longer sequences imported by interspecies transformation events tend to be much rarer, given homologous recombination’s reduced efficiency as recombinations increase in length and SNP density [32, 37]. Nevertheless, both Tn*916* and Tn*1207.1* were imported by large homologous recombinations, spanning divergent loci from other species, on multiple occasions across the species. This resolves previous genomic and experimental analyses of how these loci were acquired by pneumococci. Our results show much larger elements can be acquired by the pneumococcus via transformation from other species. Such events are clearly atypical in their properties among all detected homologous recombinations (Figure 9). Similarly the recombinations importing the *cps* loci required to escape vaccineinduced immunity are much larger than most observed around the genome (Figures 2-figure supplement 2, 3-figure supplement 2). These clinically important, but unrepresentative, recombinations do not provide evidence for the primary evolutionary benefit of transformation [112]. Rather, they likely reflect the concept underlying Milkman’s hypothesis that exchanges between divergent genotypes will only become common in the recipient where there exists an atypically strong selection pressure [113, 114]. Were they more common, genotypes would routinely converge through recombination. In these cases, the selection pressure of antibiotic consumption is sufficient to overcome the normal barriers to exchange that maintain separate streptococcal species in the human oronasopharynx [115].

The interspecies flow of sequence driven by importation of Tn*916* and Tn*1207.1* is qualitatively different to that resulting from modification of *pbp* genes. While recombination around the *pbp* genes can involve large non-pneumococcal sequence imports [58], typically the development of beta lactam resistance involved only tens or hundreds of base pair sequences exchanged within the *pbp* genes [116]. These exchanges are limited to the regions surrounding the few core genome loci directly involved in determining the resistance phenotype. However, each MGE import brings in multiple kilobases of sequence on each flank (Figure 10), and the species-wide analysis identified many different insertion sites distributed throughout the chromosome. Sequence alignments of the imported regions identify substantial structural variation in the regions surrounding the insertion site (Figures 6-figure supplement 3 - 7- figure supplement 1, 8-figure supplement 1 - 8-figure supplement 6). In this study we make no attempt to precisely determine the strain of origin for these recombination events. While our reference database is sufficient to split likely non-pneumococcal from pneumococcal DNA, it is not detailed enough to fully delineate the networks through which AMR genes spread. It is clear that *S. mitis* is a crucial source of antibiotic resistance genes for *S. pneumoniae*, the much greater diversity of *S. mitis* means the few available samples are spread thinly across the population structure [117]. Hence much greater sampling of commensal streptococcal species is needed in order to assess the most likely donor species for these interspecies transformations. This demonstrates the local sequence convergence, resulting in “fuzzy species” or “despeciation” [118,119], driven by antibiotic selection is not limited to the mosaicism of housekeeping genes associated with antibiotic resistance, but can extend throughout the streptococcal genome.

The sequence imported from the donor species can continue to permeate the recipient species in subsequent intraspecific recombinations, assuming that the resistance locus remains linked to the homologous arms on either flank. Unless preserved by selection, these flanking sequences should erode, shortening each time a recombination’s breakpoints are closer to the resistance locus than any of the previous exchanges. Hence sensitivity to detect sequence imported from the donor species will be maximal when the insertion is recent, and decline over time. Our ability to associate elements imported by transformation with their accompanying flanking sequences is further reduced by the intragenomic mobility of transposons, which may excise from their original integration site, and reinsert elsewhere in the chromosome. These are biological complications that are independent of the technical challenges of correctly inferring the co-incidence of an homologous recombination on the same phylogenetic branch, and at the same chromosomal location, as an MGE acquisition.

Some of these factors contribute to the much stronger association of Tn*1207.1* with homologous recombination events than Tn*916*. Firstly, Tn*1207.1* elements are much shorter than Tn*916*. This both makes it easier to identify the insertion site of Tn*1207.1* in contigs from draft assemblies (1,239 Tn*916* insertions are present in contigs lacking sufficient matches to a reference, as opposed to 101 for Tn*1207.1*), and makes it more likely that is can be moved by transformation, which gets exponentially less efficient at inserting elements as their length increases [37]. Secondly, Tn*916* elements were more commonly fixed in all isolates of a strain (1,072 Tn*916* insertions, compared to 41 Tn*1207.1* insertions), preventing identification of the mechanism by which the MGE integrated into the chromosome. Despite these technical challenges, Tn*916* still remained less likely to be acquired by transformation. Hence a third contributor is likely Tn*916* elements encoding machinery for transposition, including an integrase from the transposase subfamily of tyrosine recombinases. This integrase is broad in its insertion site preference, favouring sites that are AT rich or bent [20, 22]. Hence the Tn*916* element inserts at over 70 locations in the pneumococcal genome, enabling it to disassociate from any imported flanks more efficiently than Tn*1207.1*, which lacks such machinery.

Additionally, both elements can be imported by conjugation. Tn*916* encodes its own conjugative machinery, and is often found inserted into the larger Tn*5252* elements, as Tn*5253* composite elements [14, 120, 121]. Unlike Tn*916*, Tn*5252* and Tn*5253* type ICE can routinely be transferred between *S. pneumoniae* cells in vitro [37,121]. Analogously, the most commonly identified insertion site for Tn*1207.1*- type elements was within Tn*916*-type elements, sometimes themselves within a Tn*5253*-type ICE [24]. This reflects the modularity common in ICE evolution, which allows many different cargo genes to benefit from a single conjugative machinery [15, 16].

Both conjugation and transformation can allow the acquisition of resistance from other species, al- though the donor can be much less closely related in conjugation. Their relative long term contribution to resistance elements in the recipient is likely to reflect which mechanism minimises the cost of the recombinant resistant genotype. Conjugation requires the recipient host an autonomously mobile element, whereas transformation can import a resistance locus without the burden of associated MGE sequence. However, while many mobile elements have evolved to minimise the cost of their insertion site to the host cell [122], the extensive import of sequence from another species flanking the insertion is likely to be disruptive to the recipient. This is especially likely when a host gene is disrupted, as in the example of *comEC* in the German clade. Additionally, the most frequent insertion of a Tn*1207.1*-type element in the core genome was the Mega element splitting the *tag* gene, a methyladenine glycosylase involved in DNA base repair. Outside of the pneumococcus there are examples of other mobile elements inserting into mutation repair machinery, which were found to cause mutator phenotypes. For instance, in *Streptococcus agalactiae* and *Vibrio splendidus*, MGEs insert between the *mutS* and *mutL* genes involved in mismatch base repair [123–125]. These MGEs however, appear to excise during different stages of cell growth, only to reenter and disrupt the genes in question, only producing mutator phenotypes during later phases of growth. In both these species, these MGEs appear functionally under the control of the host cell [124, 125]. It is unclear if the Mega element can similarly excise and reinsert under host cell control.

In conclusion, this study has identified the broader importance of interspecies transformation in the emergence of antibiotic resistant *S. pneumoniae*. That these atypically large and SNP-dense events, originating in other streptococcal species, can be detected underscores the strong selection pressure from antibiotic consumption. This suggests there is a continual flow of sequence between related species sharing a niche, which is normally minimised by selection, but may enable rapid adaptation following public health interventions against pathogens. Even if the selection pressures are rare, transfers are sufficiently frequent for resistant genotypes to emerge and spread locally, but particularly successful genotypes, such as PMEN3, can rapidly spread between continents. This highlights the challenges of blocking the transfer of resistance loci into pathogenic species.

## Supplementary Materials

**Figure 2-figure supplement 1:**
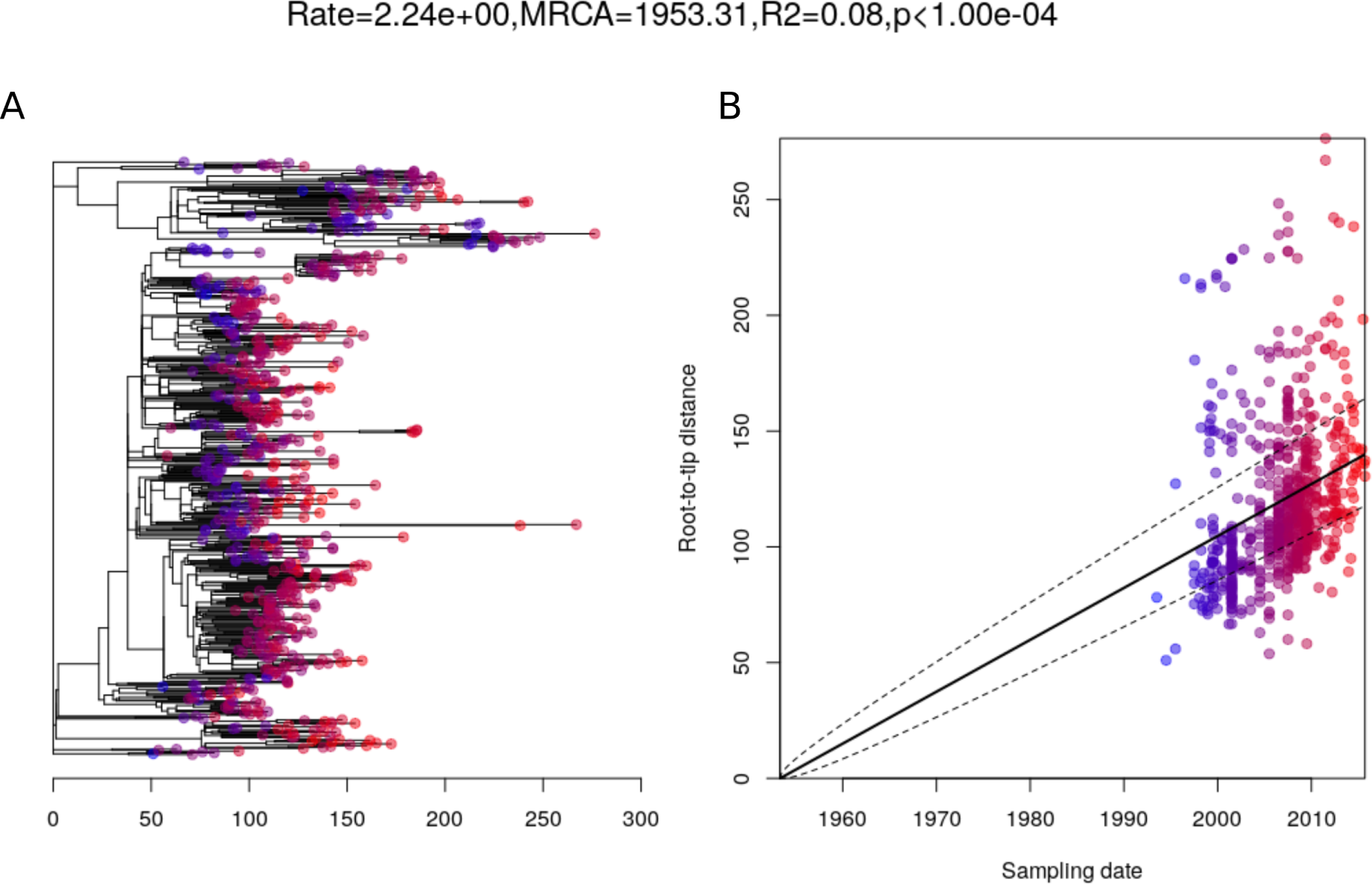
Root to tip analysis of PMEN3 Lineage. A. Represents the 663 isolate phylogeny with node tips coloured by date of isolation. **B** Linear regression of root to tip distance against sampling date for Isolates.

**Figure 2-figure supplement 2:**
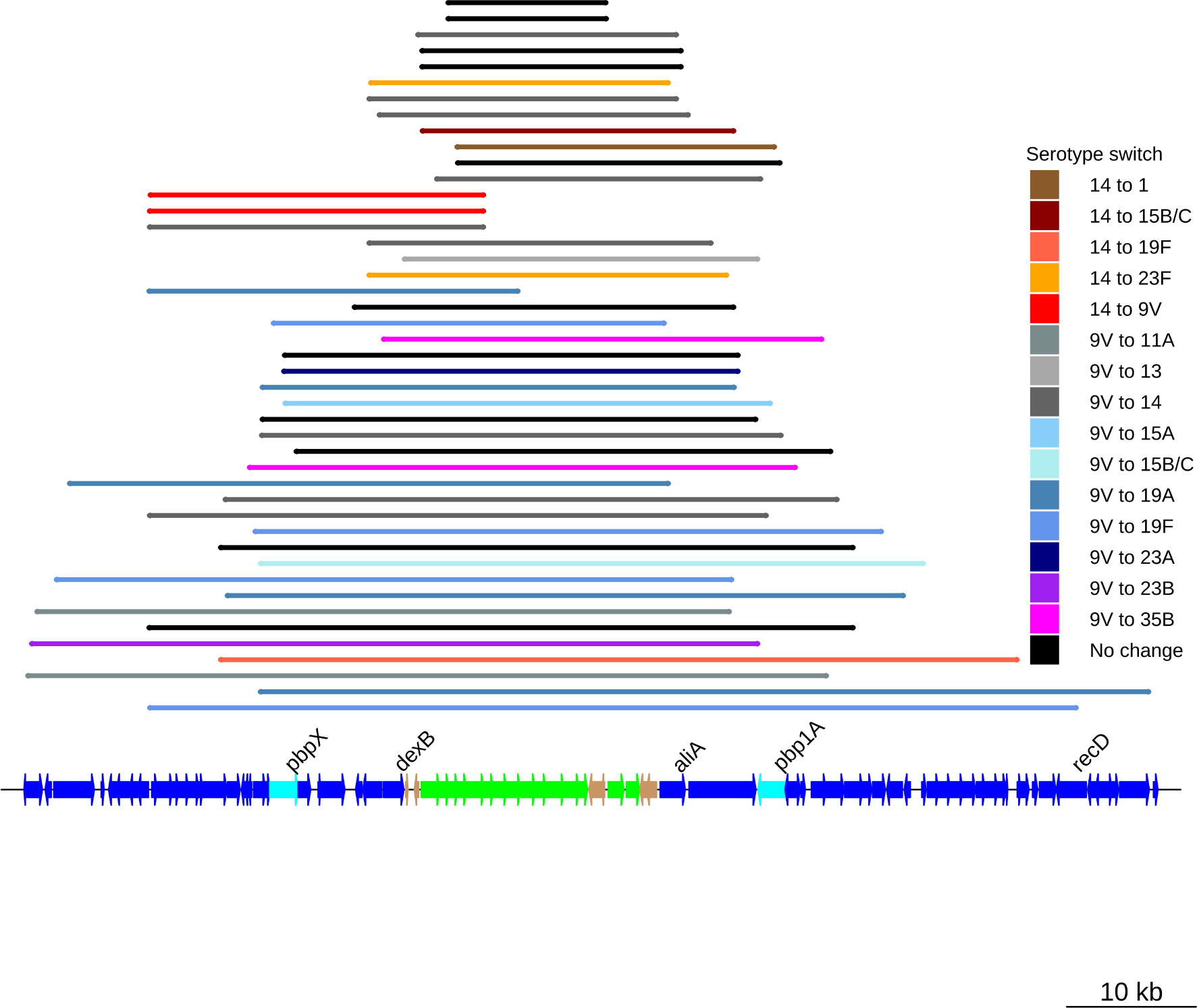
Serotype switching events across the PMEN3 collection. Coloured bars represent the recombination events associated with these switches in serotype. The bars map to the genome annotation below, with the length of the bar indicating a longer recombination event. The *cps* and surrounding loci are highlighted below, with some recombination events spanning further across the *pbp1a* and *pbp2x* genes as well. The black bars represent recombinations across the *cps* loci that weren’t associated with a serotype switch.

**Figure 3-figure supplement 1:**
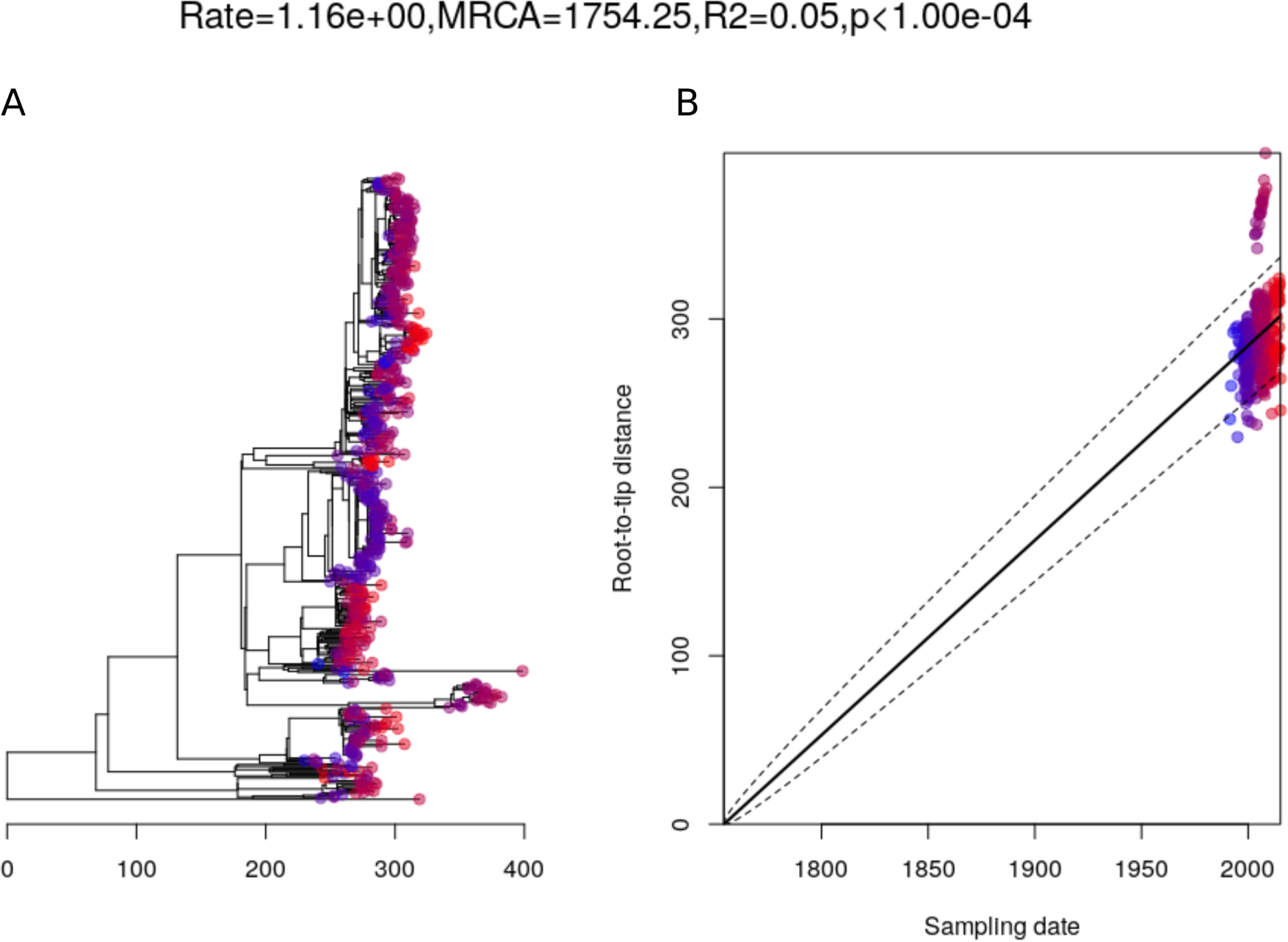
Root to tip analysis of PMEN9 Lineage. A. Represents the trimmed 529 isolate phylogeny with node tips coloured by date of isolation. **B** Linear regression of root to tip distance against sampling date for Isolates.

**Figure 3-figure supplement 2:**
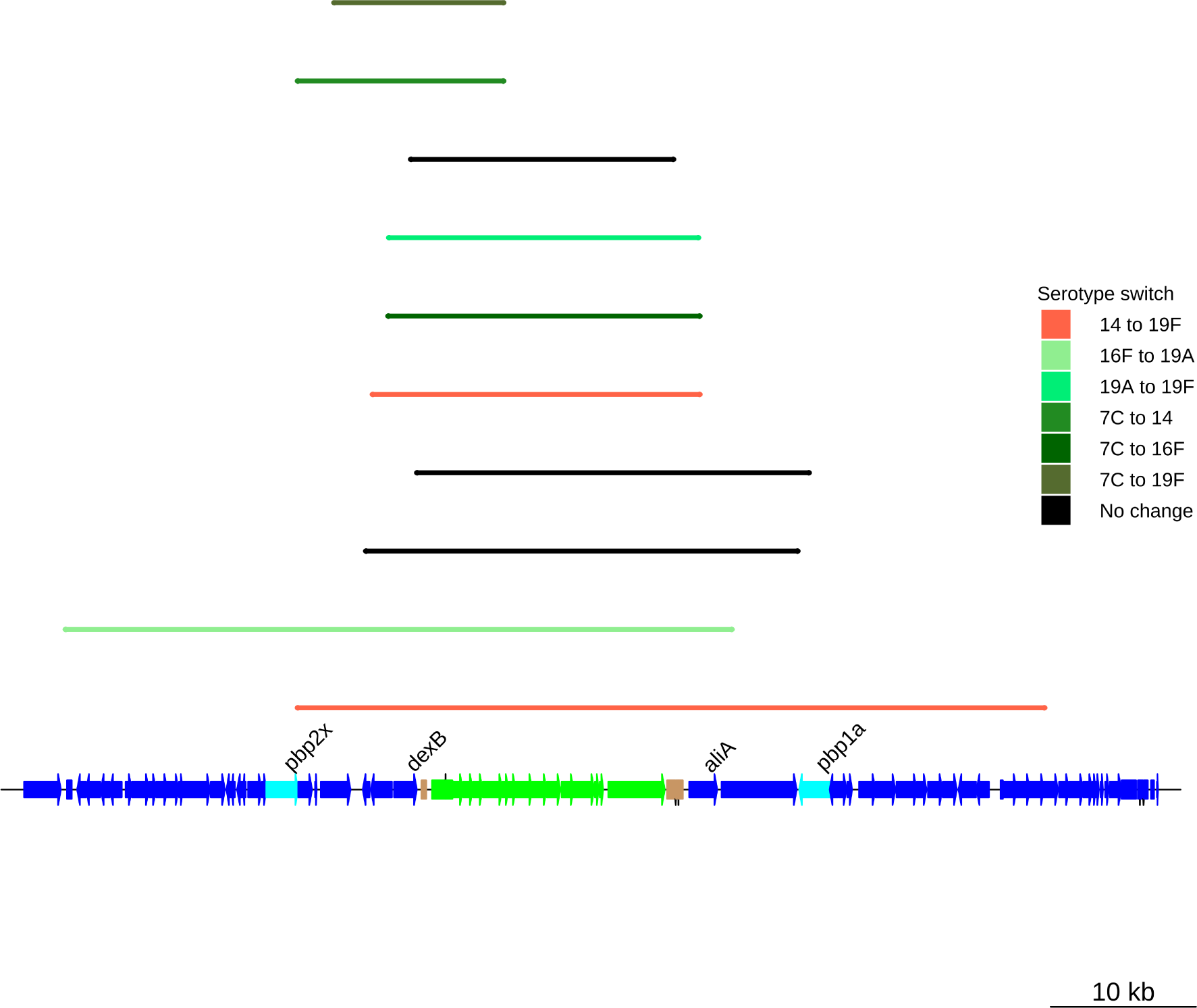
Serotype switching events across the PMEN9 collection. Coloured bars represent the recombination events associated with these switches in serotype. The bars map to the genome annotation below, with the length of the bar indicating a longer recombination event. The *cps* and surrounding loci are highlighted below, with some recombination events spanning further across the *pbp1a* and *pbp2x* genes as well. The black bars represent recombinations across the *cps* loci that weren’t associated with a serotype switch.

**Figure 5-figure supplement 1:**
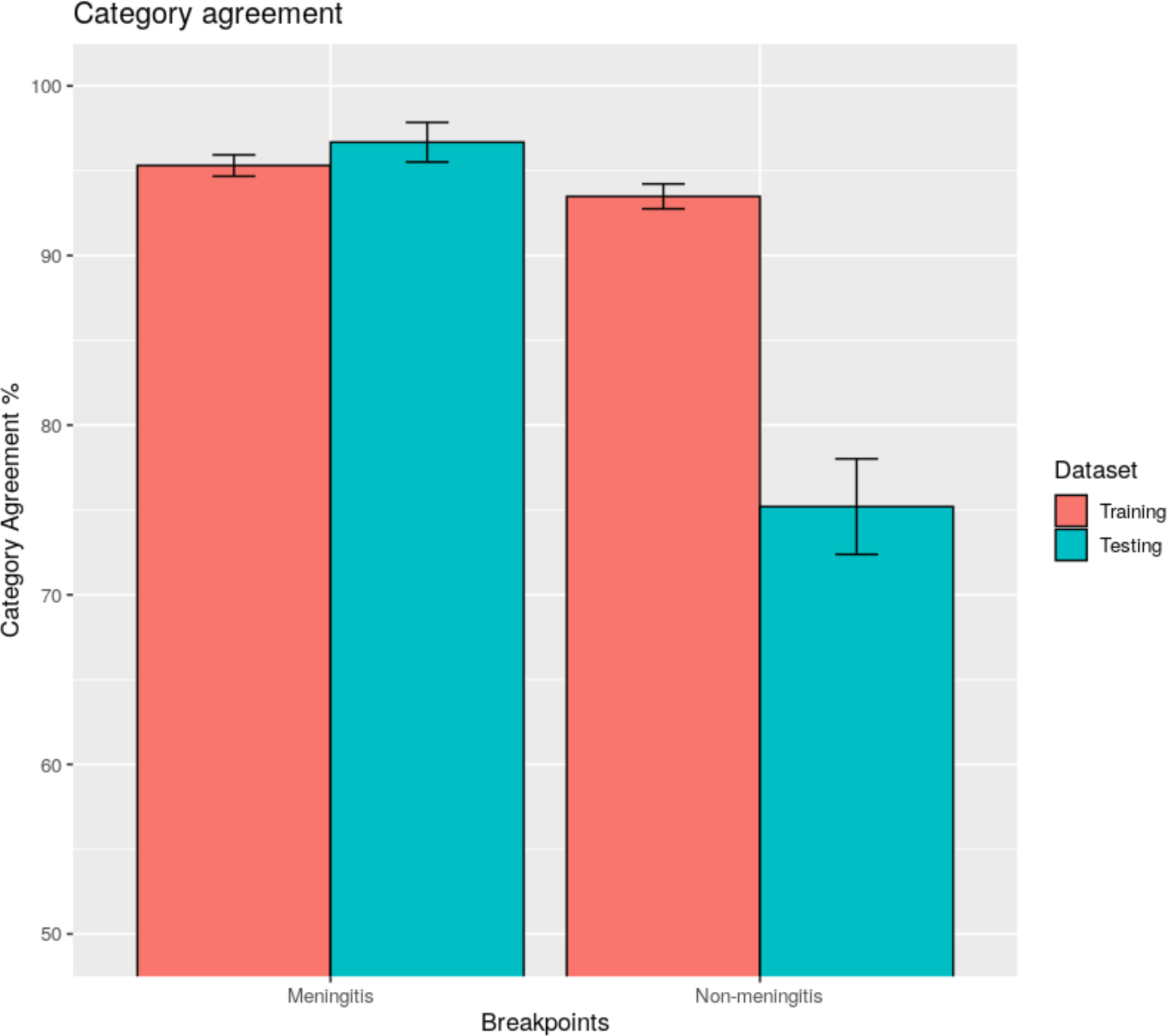
Comparisons of Category Agreement across different penicillin resistance breakpoints. Bars represent the category agreement, the number of categories correctly predicted across the testing or training dataset, for the two breakpoint sets tested. The training data for the model corresponds to the 4,342 isolate CDC datset, while the testing dataset refers to the 903 PMEN lineage isolates with available MIC data.

**Figure 5-figure supplement 2:**
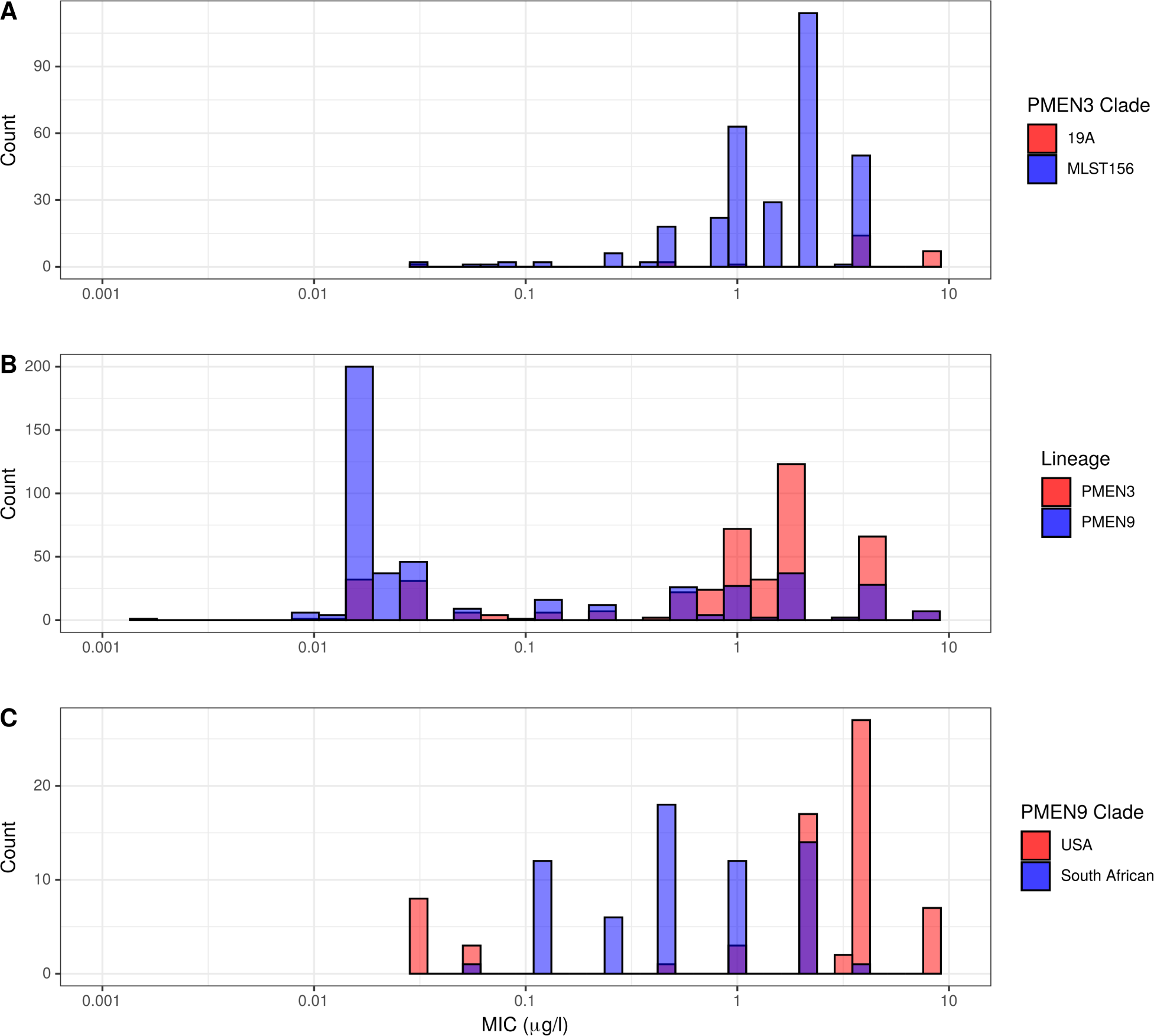
Histograms of recorded MIC values for penicillin across the PMEN3 and PMEN9 collections. A. MIC values for penicillin for the primarily resistant clades ST156 and 19A within the PMEN3 collection. N = 313 for the ST156 clade and N = 25 for the 19A clade. **B** MIC for penicillin across both PMEN3 and PMEN9. N = 438 for PMEN3 and N = 465 for PMEN9. **C** MIC values for penicillin for the primarily resistant USA and South African clades within the PMEN9 collection. N = 68 for the USA clade and N = 64 for the South African clade.

**Figure 5-figure supplement 3:**
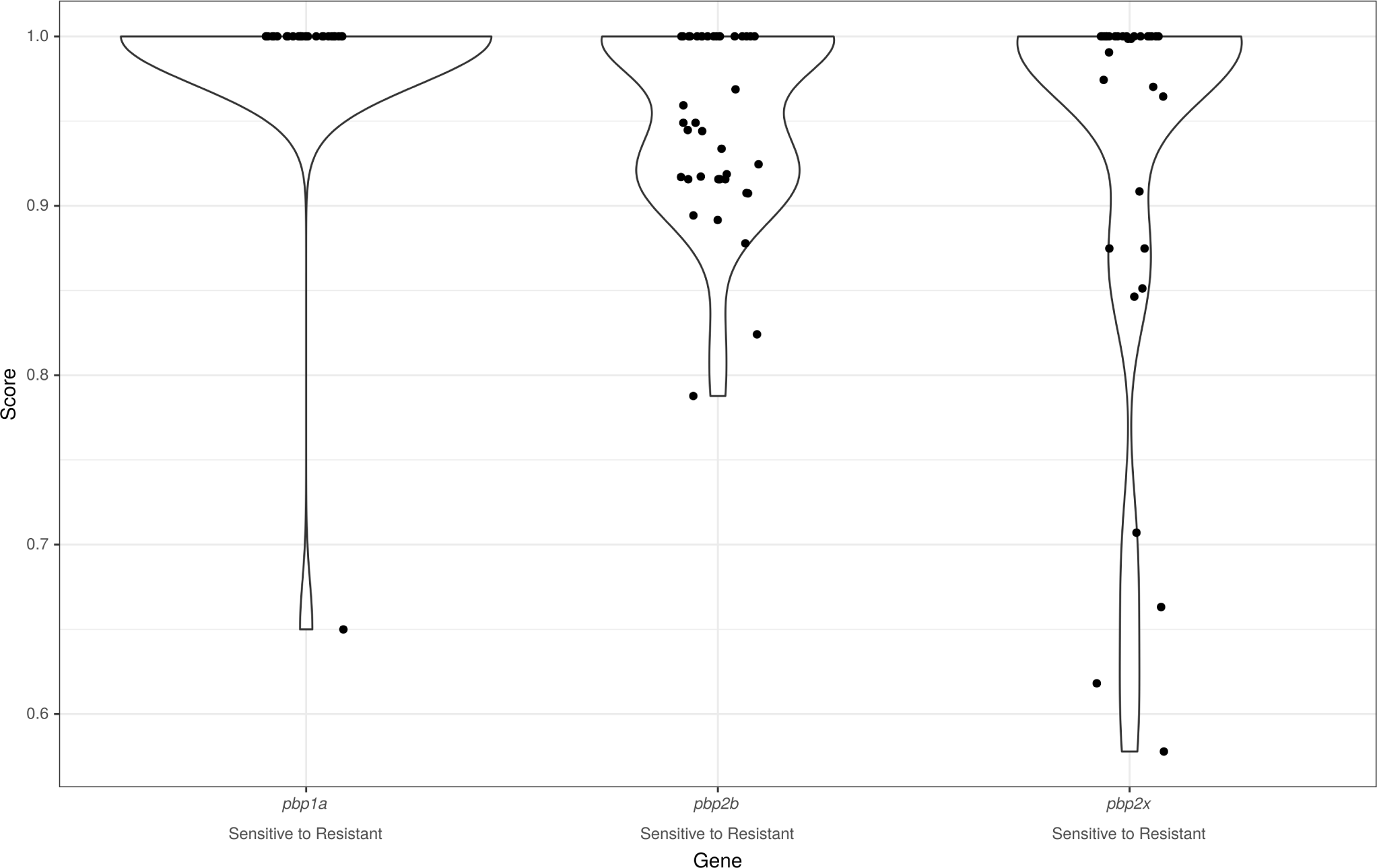
Origin of *pbp* genes for penicillin resistant isolates. Graph depicts the distribution of *γ* scores for *pbp* gene sequences within the gps collection where these genes are present within a gain of resistance associated homologous recombination event. Dots represent individual genes scores within recombination events. The wider distribution or scores is summarised by the violin plot outline.

**Figure 5-figure supplement 4:**
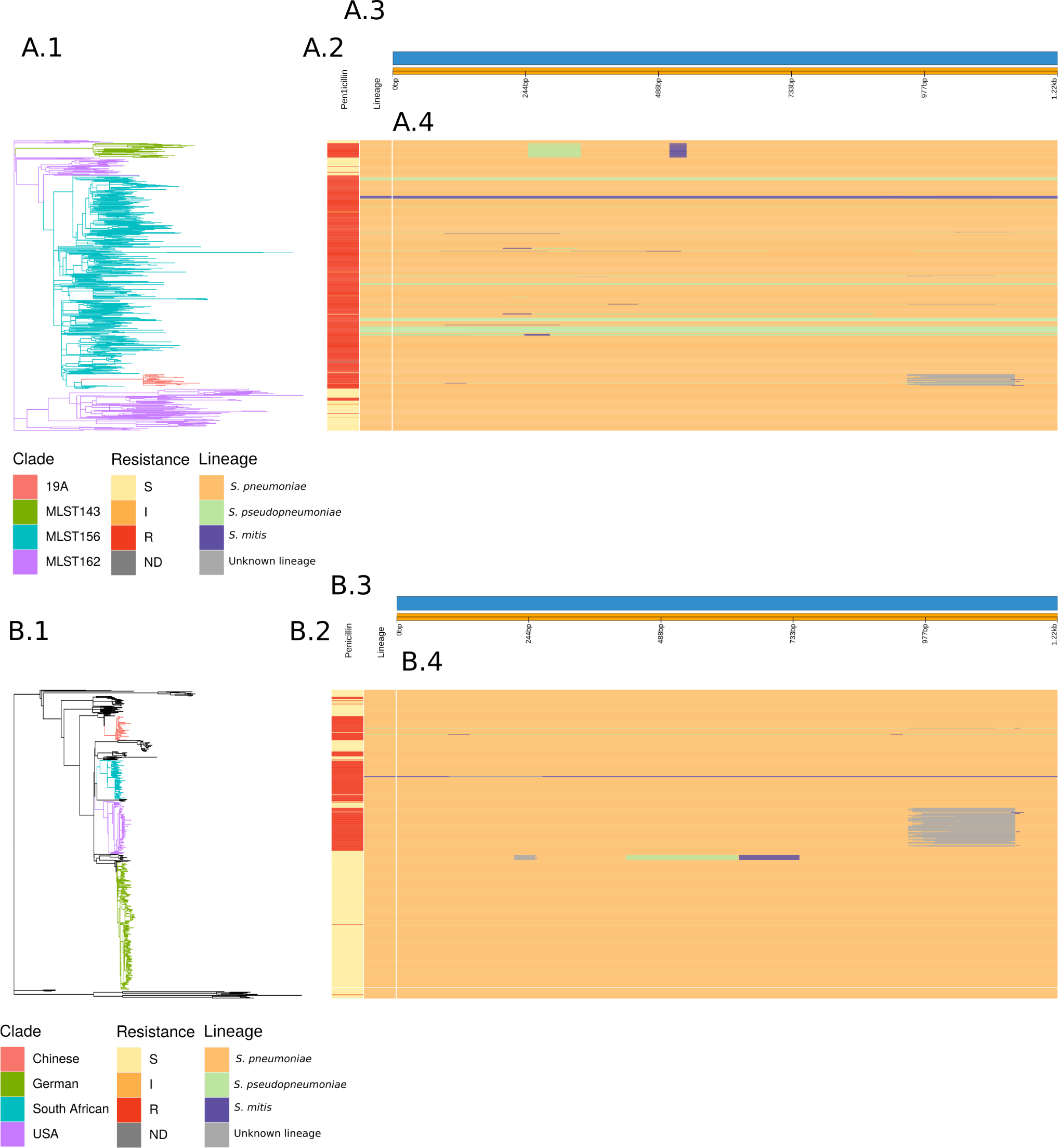
Analysis of the origin of the *murM* gene across the PMEN3 and PMEN9 Lineages. A.1. The Recombination-corrected whole genome phylogeny of PMEN3, with branches coloured by clade of interest. **A.2** Bars indicating the penicillin resistance category of isolates and the lineage inferred for the *murM* gene of each isolate. **A.3** Representation of the *murM* gene. **A.4** This panel shows the inferred lineage for each base of each sequence across the PMEN3 phylogeny. Solid horizontal bars indicate sequences belonging to a particular lineage, as indicated by the colour. Changes in colour across a bar indicate different *murM* segments were inferred to originate in different lineages, suggesting a mosaic allele generated by recombination. **B.1** Recombination-corrected whole genome phylogeny of PMEN9, with branches coloured by clade of interest. **B.2** Bars indicating the penicillin resistance category of isolates and the overall lineage inferred for the *murM* gene of each isolate. **B.3** Representation of the *murM* gene. **B.4** This panel shows the inferred lineage for each base of each sequence across the PMEN9 phylogeny, as for the upper part of the figure.

**Figure 6-figure supplement 1:**
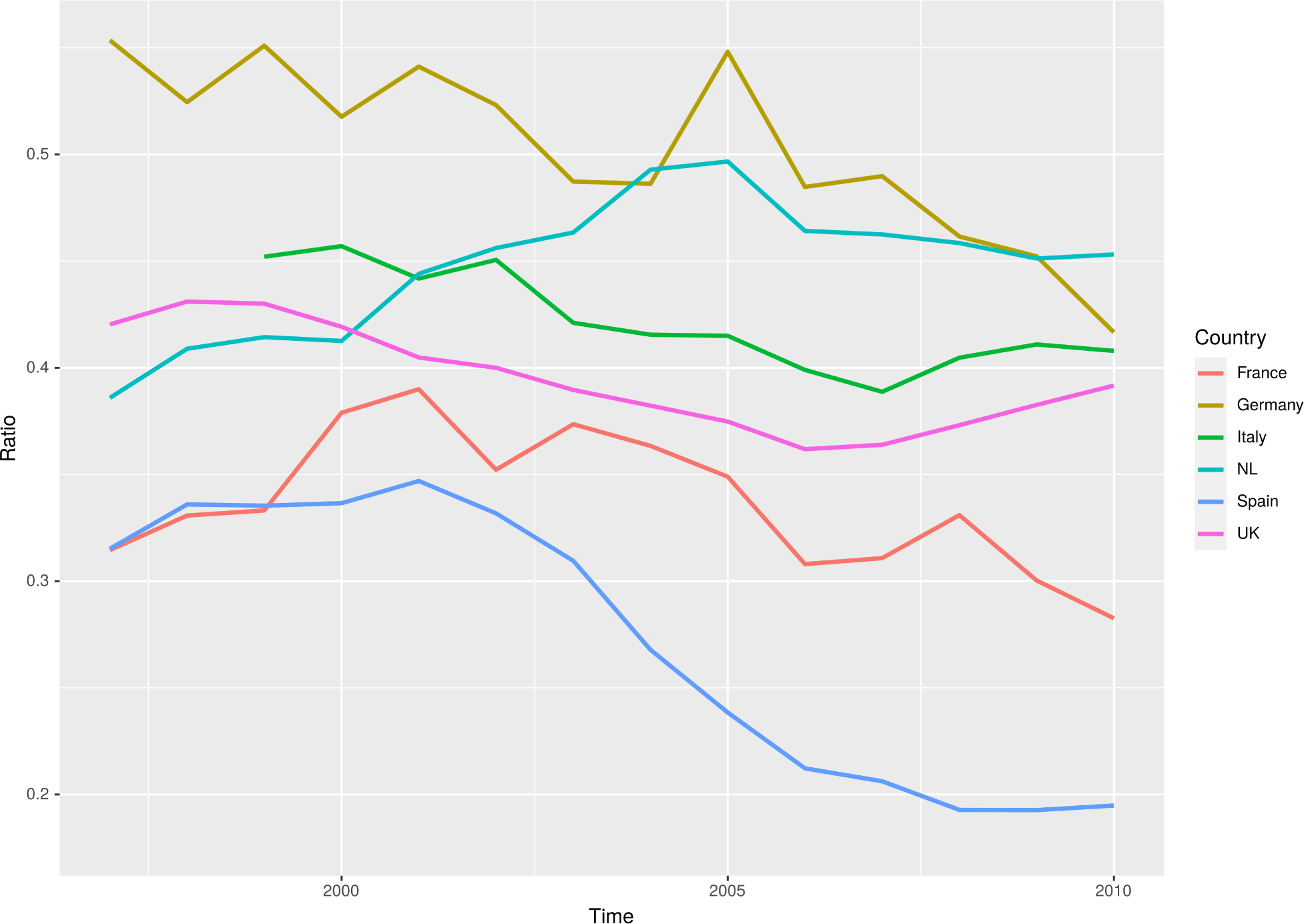
Ratio of macrolide to penicillin consumption in Europe. The ratios of macrolide use, in DDD, per penicillin use for 6 major European countries across a 13 year period from 1997 to 2010.

**Figure 6-figure supplement 2:**
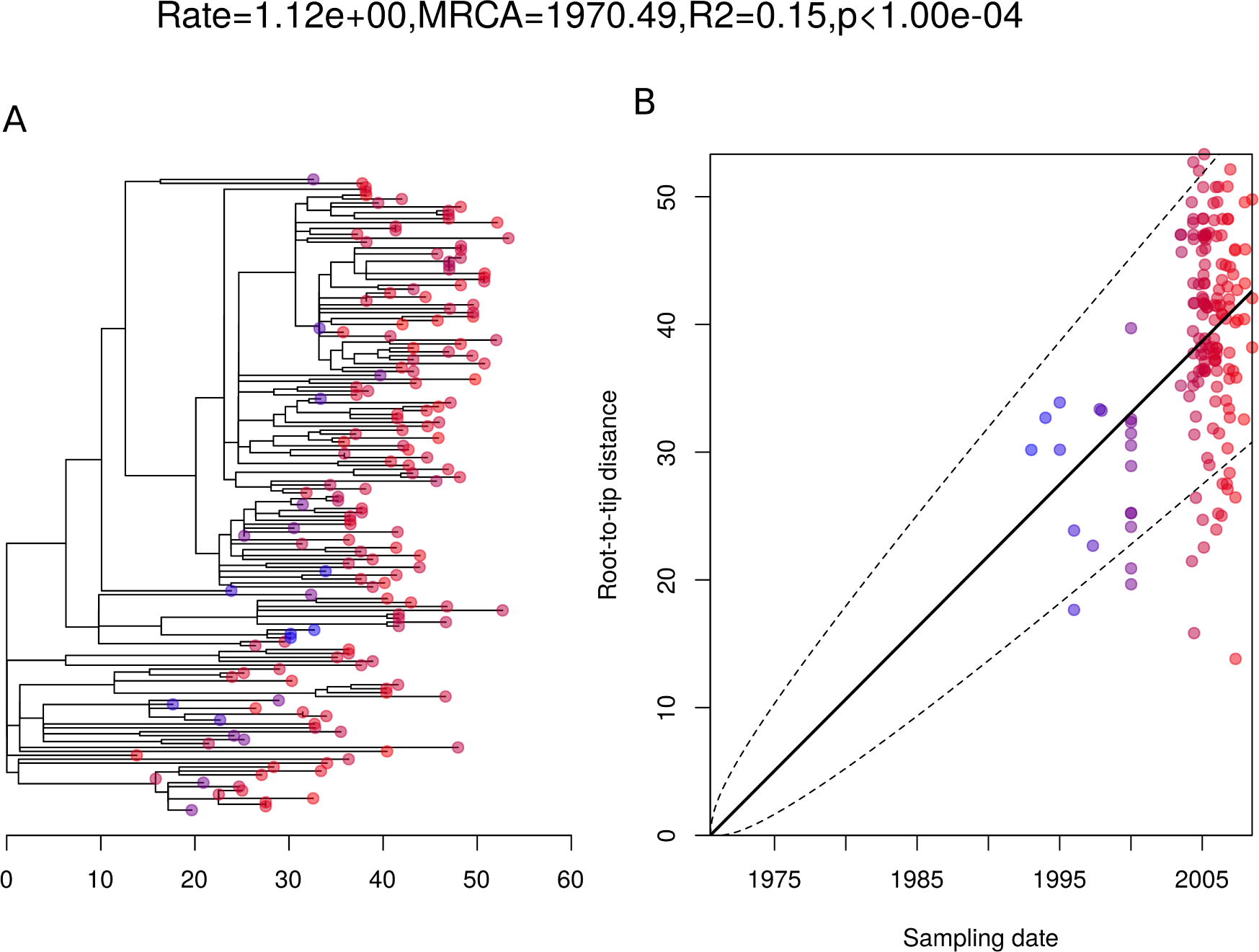
Root to tip analysis of 162 German isolates within PMEN9. A. Represents the 162 isolate phylogeny with node tips coloured by date of isolation. **B** Linear regression of root to tip distance against sampling date for Isolates.

**Figure 6-figure supplement 3:**
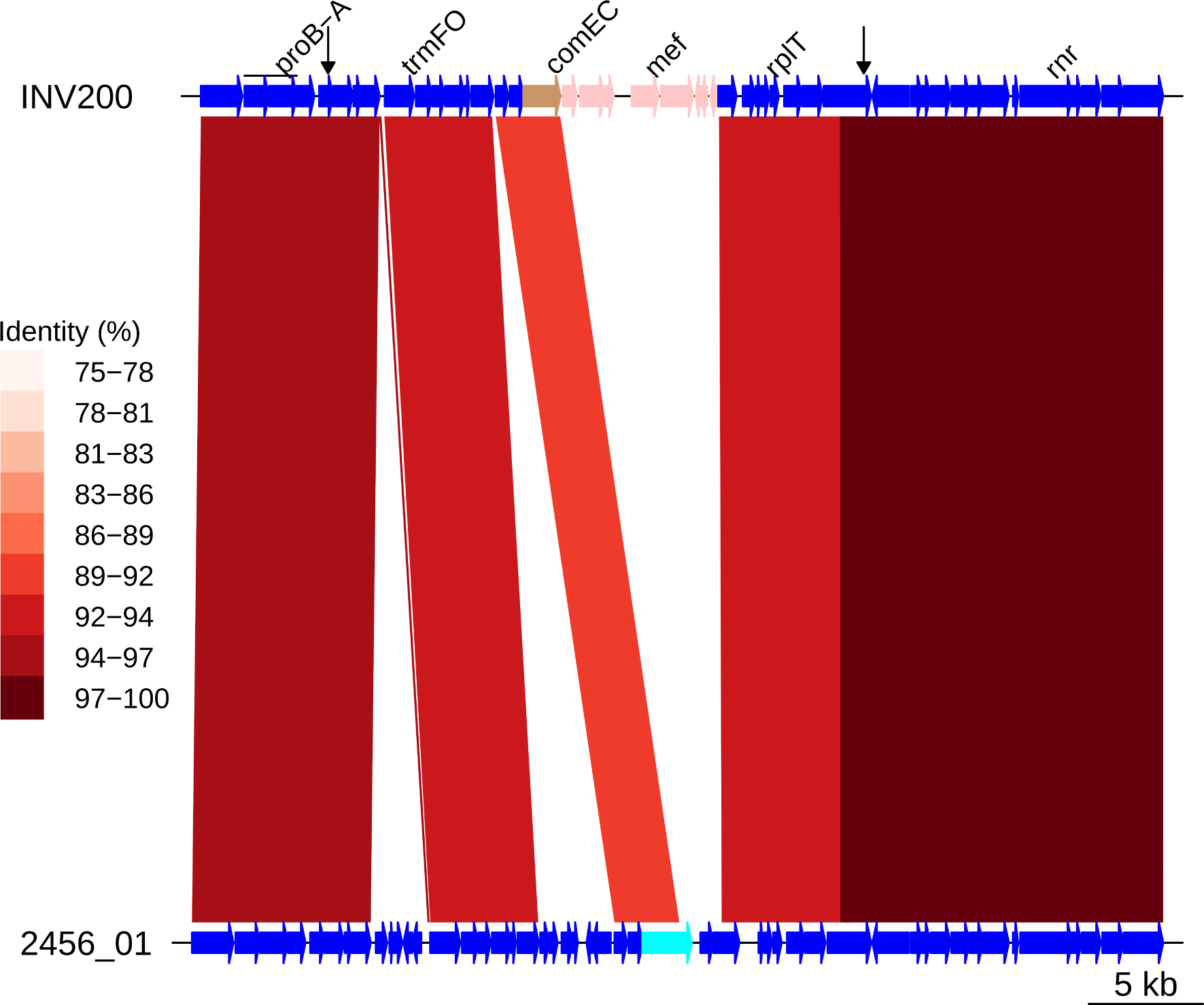
Insert of Tn*1207.1* within PMEN9 reference genome. Comparison of the Tn*1207.1* element insertion, highlighted in pink within the INV200 genome, and the 2456_01 sample with an intact *comEC*. The split *comEC* gene is highlighted in brown within INV200, cyan within 2456_01. Bars between the genome represent sequence matches, with these bars shaded by % identity between the sequences. Arrows along the INV200 genome mark the start and end of the recombination event bringing in the Tn*1207.1* element.

**Figure 7-figure supplement 1:**
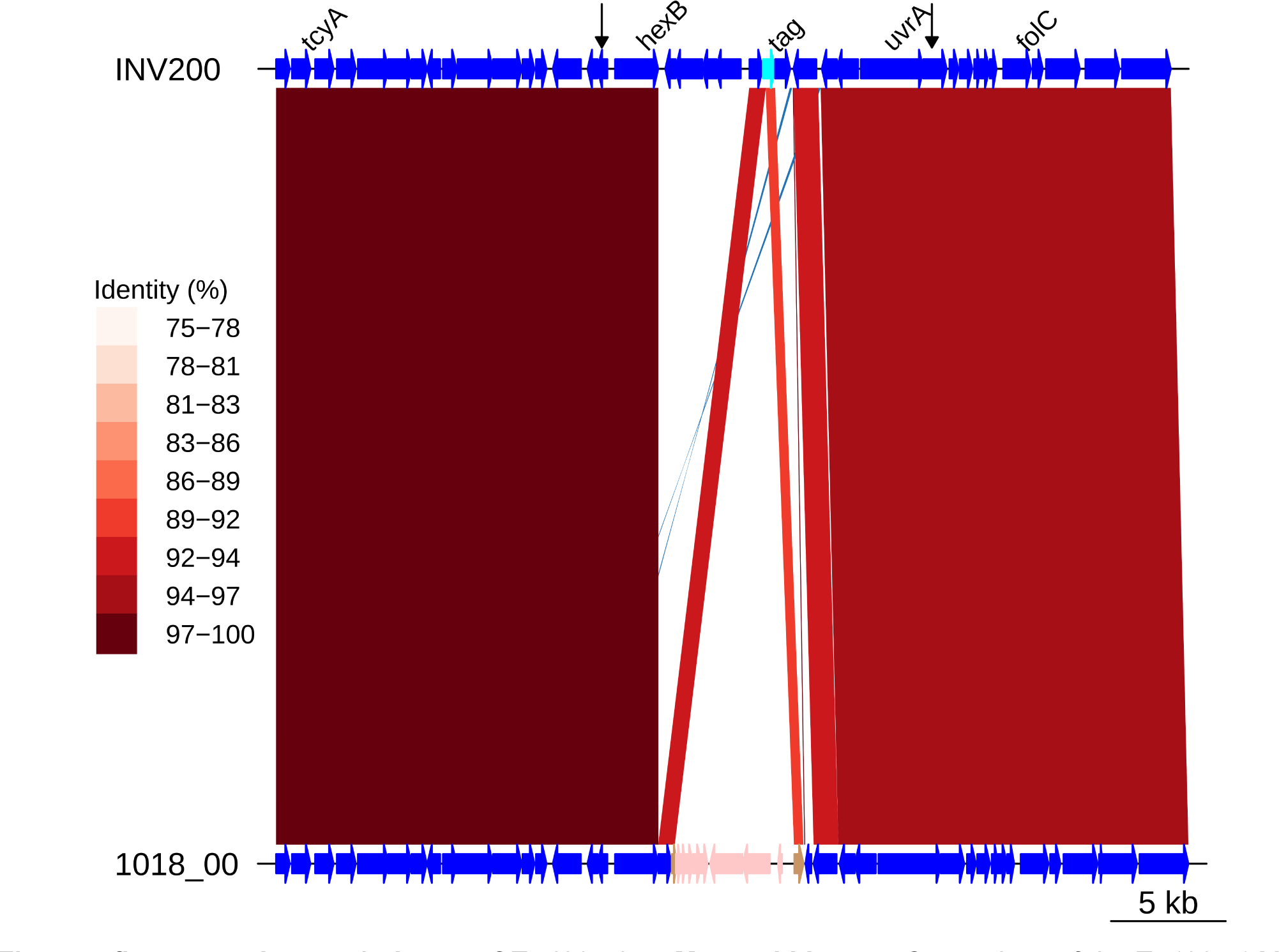
Insert of Tn*1207.1* as Mega within *tag*. Comparison of the Tn*1207.1* Mega element insertion, highlighted in pink within the 1018_00 genome, and the INV200 sample with an intact *tag*. The split *tag* gene is highlighted in brown within 1018_00 and the complete gene in cyan within INV200. Bars between the genome represent sequence matches, with these bars shaded by % identity between the sequences. Arrows along the 1018_00 genome mark the start and end of the recombination event bringing in the Tn*1207.1* Mega element.

**Figure 10-figure supplement 1:**
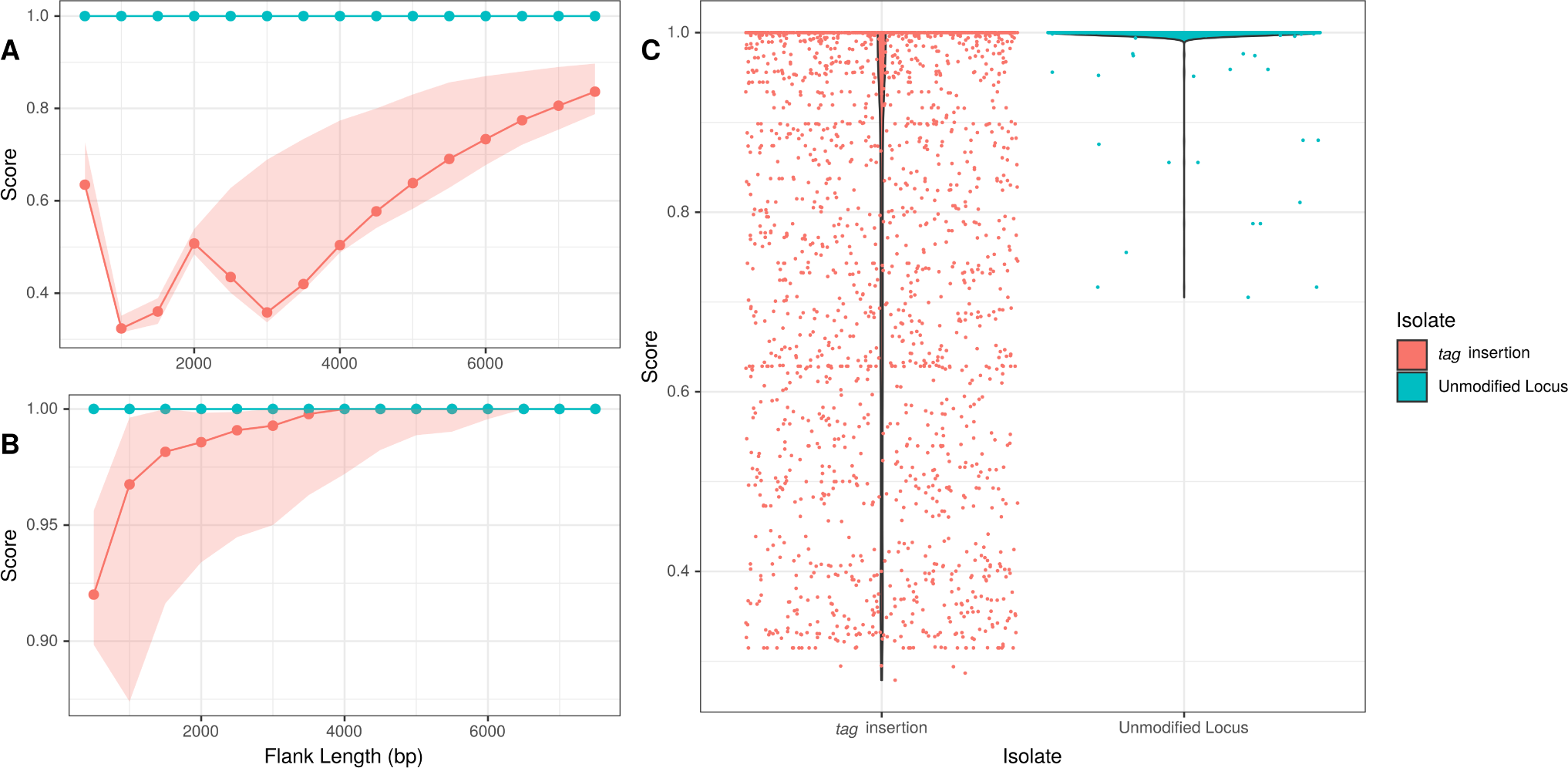
Flanking region origin for Tn*1207.1 tag* insertions. A. The median *γ* score of upstream flanking regions of the Tn*1207.1* tag insertion insertion events. The median *γ* score for the homolgous regions in isolates without the MGE, and hence with an unmodified locus, are highlighted in cyan. Shaded regions represent the Inter-quartile range (IQR) of the *γ* score. **B** The *γ* score for regions extracted downstream of the insertion. Shaded regions represent the IQR of the *γ* score. **C** The distribution of *γ* scores across flanking lengths for flanks upstream and downstream of the Tn*1207.1 tag* inserts, for both isolates with the insert and those with an unmodified homologous region.

**Figure 8-figure supplement 1:**
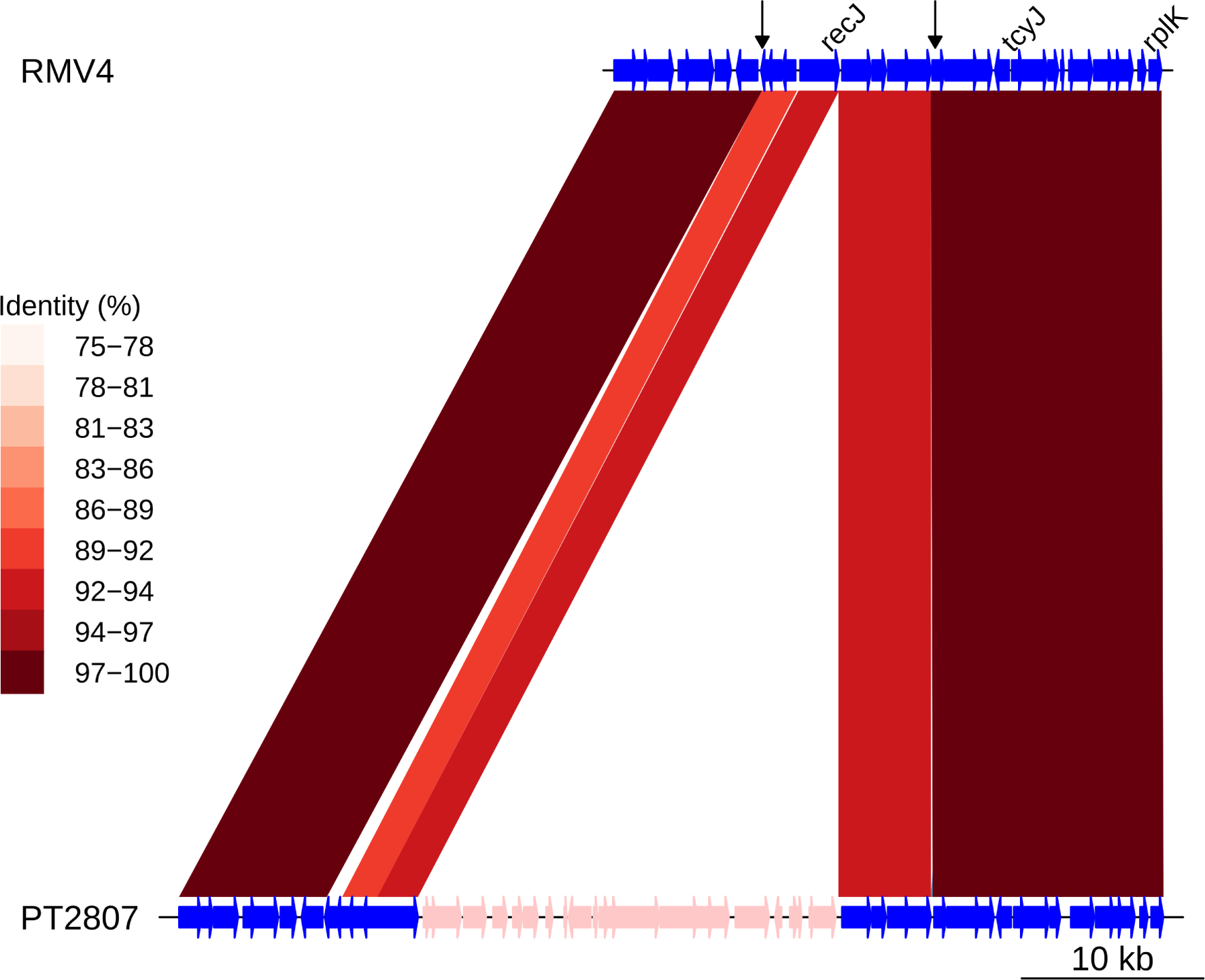
**Insert of Tn*916* downstream of *recJ***. Comparison of the Tn*916* element insertion, highlighted in pink within the PT2807 genome, and the RMV4 sample with no insertion. Bars between the genome represent sequence matches, with these bars shaded by % identity between the sequences. Arrows along the RMV4 genome mark the start and end of the recombination event bringing in the Tn*916* element.

**Figure 8-figure supplement 2:**
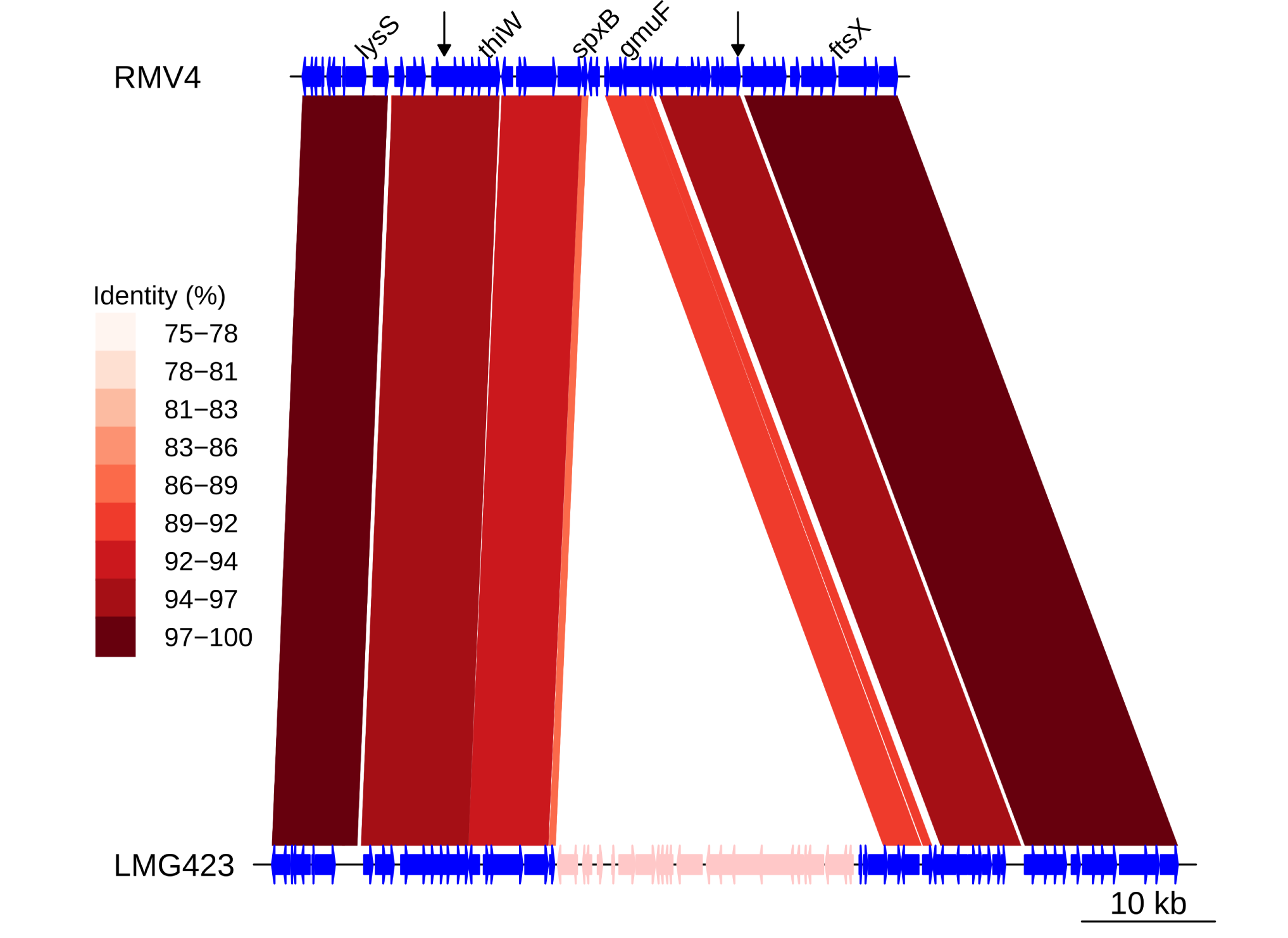
**Insert of Tn*916* downstream of *gmuF*** . Comparison of the Tn*916* element insertion, highlighted in pink within the LMG423 genome, and the RMV4 sample with no insertion. Bars between the genome represent sequence matches, with these bars shaded by % identity between the sequences. Arrows along the RMV4 genome mark the start and end of the recombination event bringing in the Tn*916* element.

**Figure 8-figure supplement 3:**
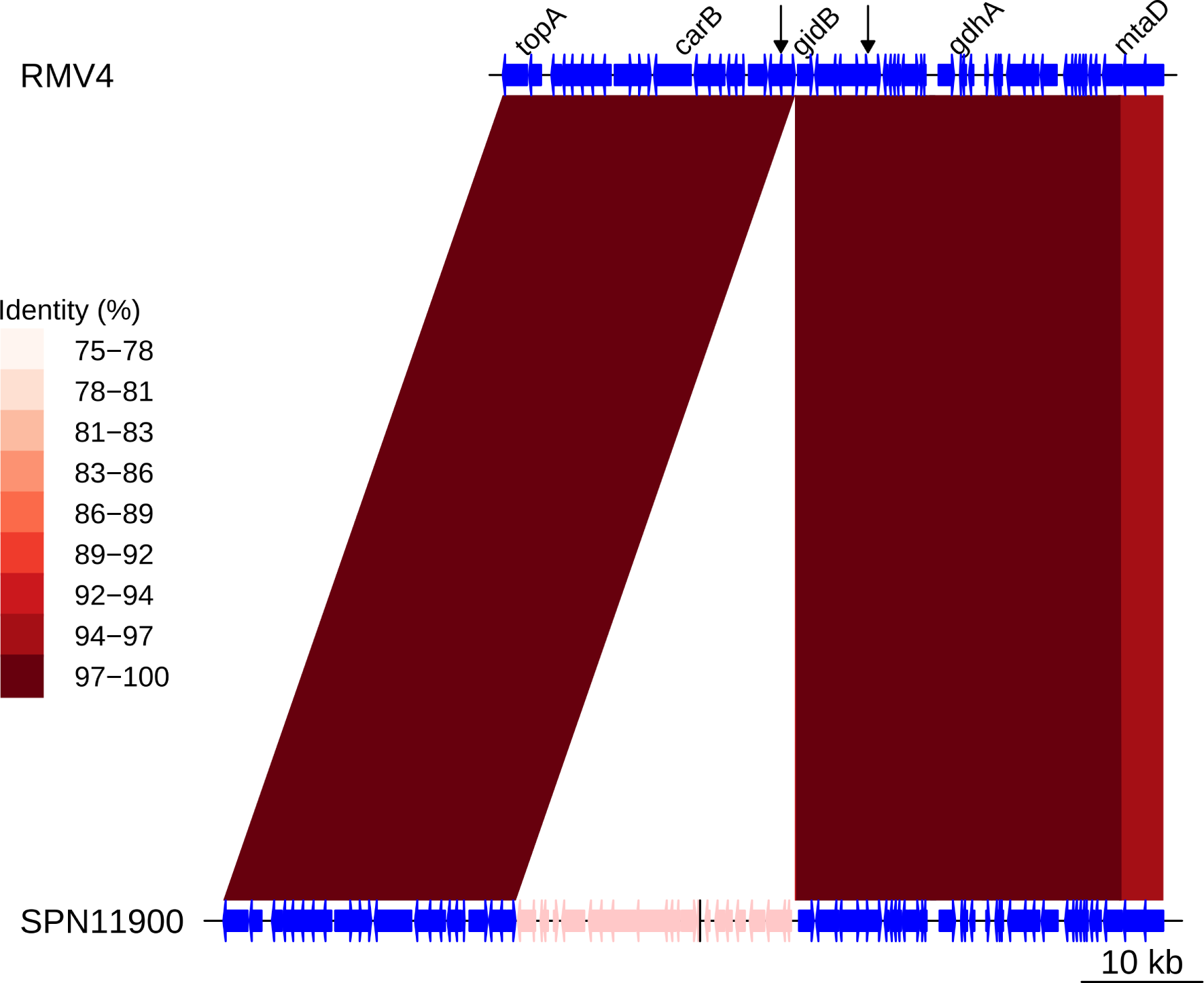
**Insert of Tn*916* upstream of *gidB***. Comparison of the Tn*916* element insertion, highlighted in pink within the SPN11900 genome, and the RMV4 sample with no insertion. Bars between the genome represent sequence matches, with these bars shaded by % identity between the sequences. Arrows along the RMV4 genome mark the start and end of the recombination event bringing in the Tn*916* element.

**Figure 8-figure supplement 4:**
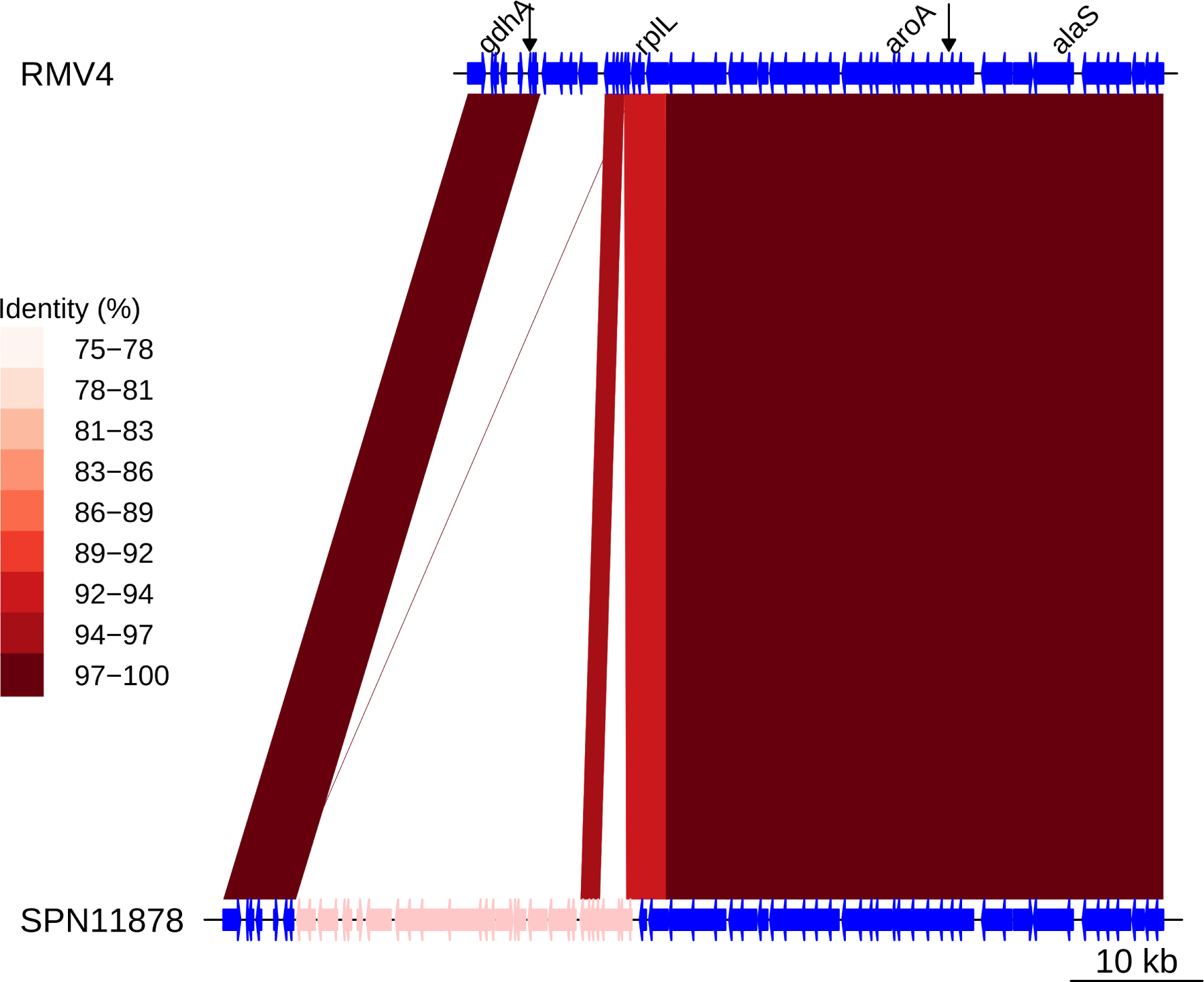
**Insert of Tn*916* upstream of *rplL***. Comparison of the Tn*916* element insertion, highlighted in pink within the SPN11878 genome, and the RMV4 sample with no insertion. Bars between the genome represent sequence matches, with these bars shaded by % identity between the sequences. Arrows along the RMV4 genome mark the start and end of the recombination event bringing in the Tn*916* element.

**Figure 8-figure supplement 5:**
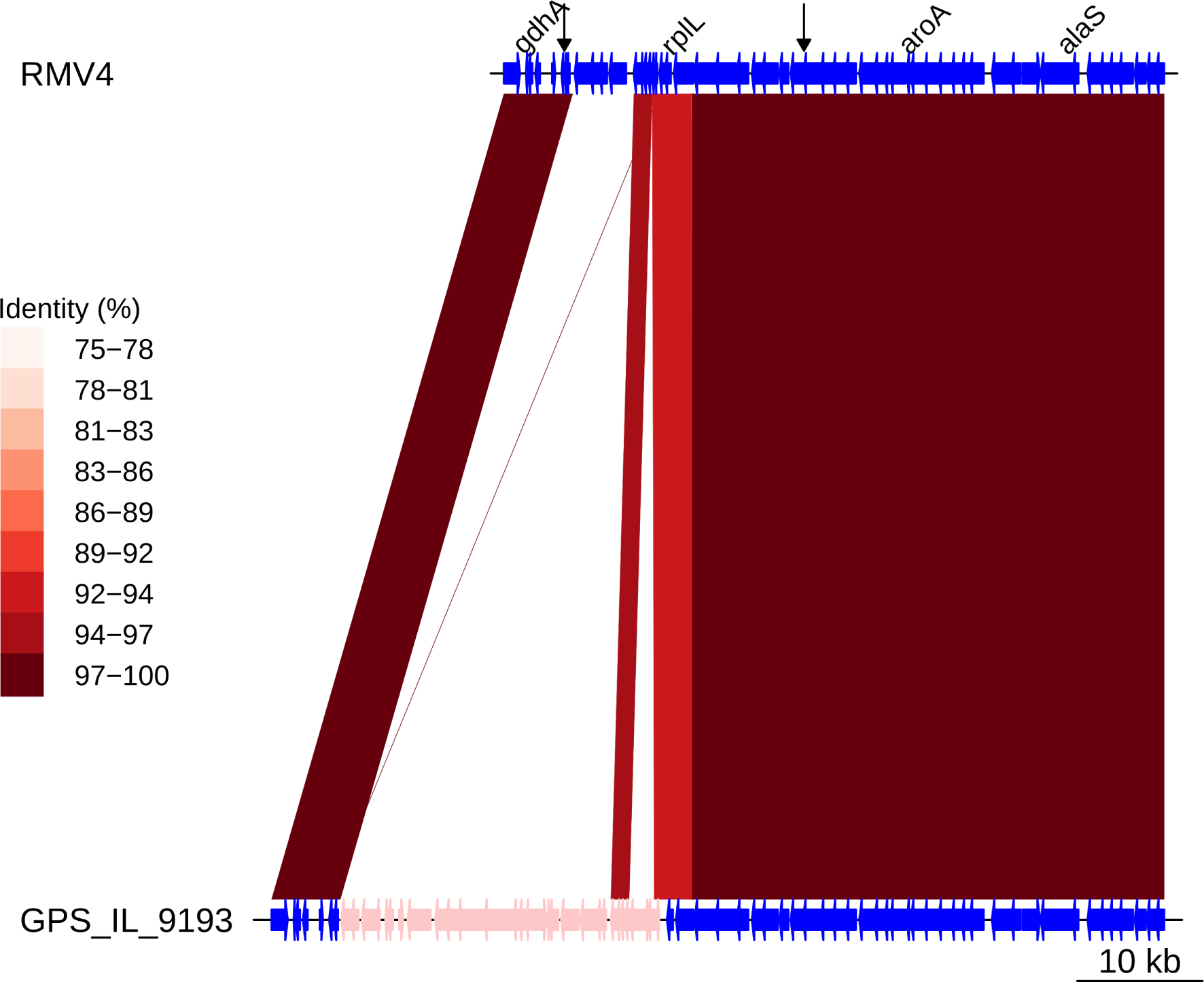
**Insert of Tn*916* upstream of *rplL***. Comparison of the Tn*916* element insertion, highlighted in pink within the GPS_IL_9193 genome, and the RMV4 sample with no insertion. Bars between the genome represent sequence matches, with these bars shaded by % identity between the sequences. Arrows along the RMV4 genome mark the start and end of the recombination event bringing in the Tn*916* element.

**Figure 8-figure supplement 6:**
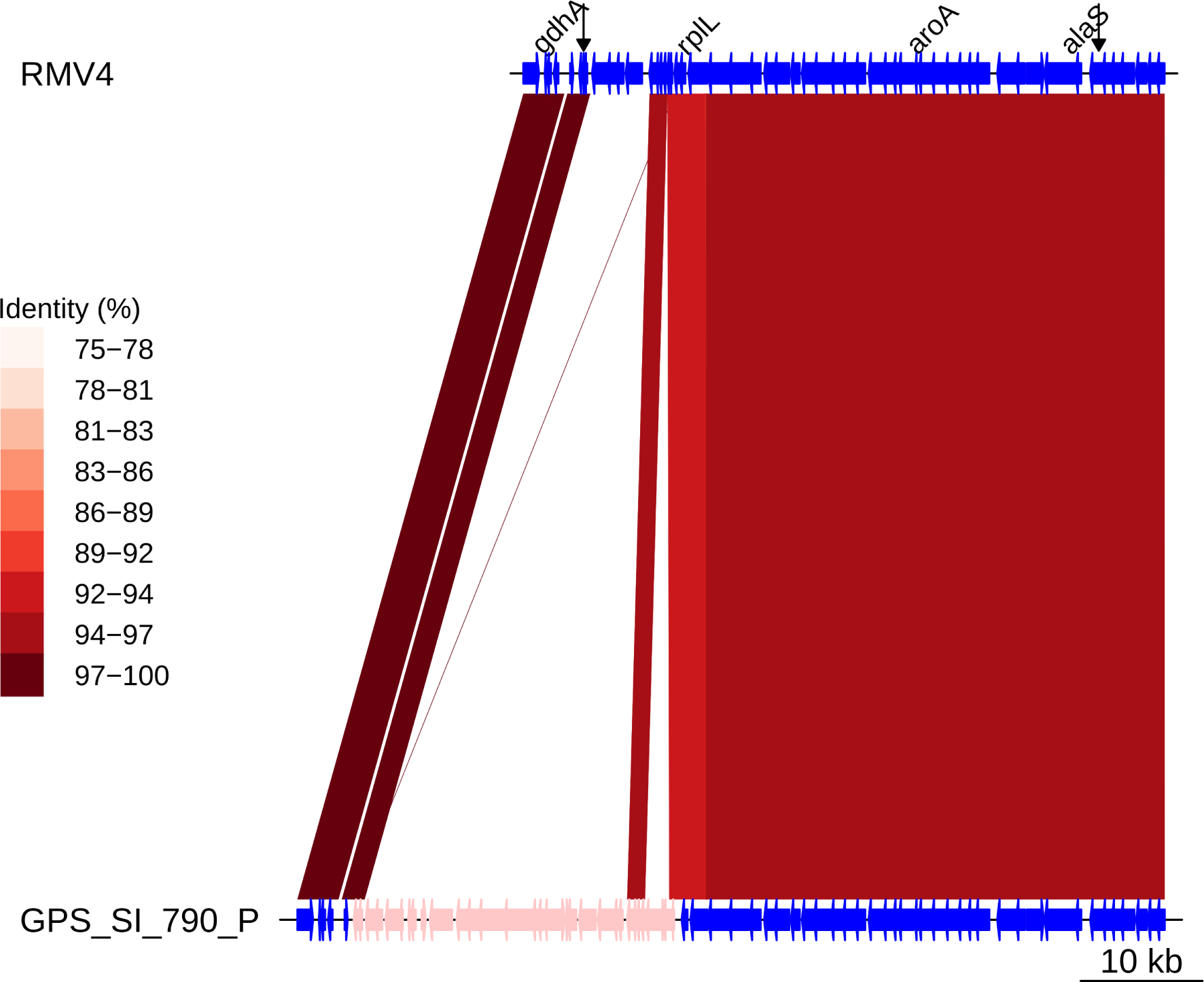
**Insert of Tn*916* upstream of *rplL***. Comparison of the Tn*916* element insertion, highlighted in pink within the GPS_SI_790_P genome, and the RMV4 sample with no insertion. Bars between the genome represent sequence matches, with these bars shaded by % identity between the sequences. Arrows along the RMV4 genome mark the start and end of the recombination event bringing in the Tn*916* element.

